# High-coverage genome sequencing of Yayoi and Jomon individuals shed light on prehistoric human population history in East Eurasian

**DOI:** 10.1101/2024.08.09.606917

**Authors:** Koji Ishiya, Fuzuki Mizuno, Jun Gojobori, Masahiko Kumagai, Yasuhiro Taniguchi, Osamu Kondo, Masami Matsushita, Takayuki Matsushita, Li Wang, Kunihiko Kurosaki, Shintaroh Ueda

**Affiliations:** Department of Biological Sciences, Graduate School of Science, The University of Tokyo, Tokyo, Japan; Kanazawa University, Kanazawa, Japan; Department of Legal Medicine, Toho University School of Medicine, Tokyo, Japan; Department of Evolutionary Studies of Biosystems, SOKENDAI (The Graduate University for Advanced Studies), Hayama, Japan; Advanced Analysis Center, National Agriculture and Food Research Organization, Tsukuba, Japan; Department of Archaeology, Faculty of Letters, Kokugakuin University, Tokyo, Japan; The Doigahama Site Anthropological Museum, Yamaguchi, Japan

## Abstract

The migration of prehistoric humans led to intriguing interactions and changes in cultural and genetic heritage. In Eurasia, prehistoric migration and population replacement have affected present-day humans. The available high-quality genetic evidence for prehistoric migration in eastern Eurasia, particularly in the Far East, is still limited. We succeeded in obtaining low-contaminant, high-coverage genomes from middle-Yayoi (>46-fold coverage) and Initial Jomon (>67-fold coverage) individuals from mainland Japan. This study demonstrated that the Yayoi individual exhibited a genetic profile distinct from that of the indigenous Jomon population of the Japanese archipelago, suggesting that Yayoi ancestry was connected to the peopling of the Eurasian continent. Our high-coverage genome provides interesting insights into the evolution of copy number polymorphisms related to the dietary styles of ancient Japanese people. The copy number estimates of the amylase gene for the Yayoi individual were comparable to those of present-day East Asians who have diets high in starch. This suggests that the population in the middle Yayoi period may have already adapted to high-starch diets, which may have been related to paddy rice agriculture introduced from the continent. Furthermore, the individual from the initial Jomon period showed high amylase copy numbers comparable to those from modern East Eurasia, including modern Japanese. This suggests that some Jomon people may have consumed a high-starch diet then. The high-coverage whole-genome sequence also revealed differences in the demographic backgrounds of the two ancestral populations during the Yayoi and Jomon periods. Our results shed light on the prehistorical events and origins of related migrations from Eurasia at that time and their genetic background, cultural transformations, and links to modern Japanese people.

## 1. Introduction

The dispersal events of anatomically modern humans into Eurasia for approximately 60,000 years have been posited in the traditional out-of-Africa model. However, their migration routes and peopling history remain controversial in Eastern Eurasia (Bae et al., 2017; Matsumura et al., 2018). Prehistoric genetic evidence of population replacement and admixture can shed light on the people in these regions. Recent genetic studies of ancient humans in East Asia have provided novel insights into the events of dispersal and population replacement since the Late Pleistocene in the Eurasian continent (Ning et al., 2020; Yang et al., 2020; Gakuhari et al., 2020; Wang et al., 2021; Cooke et al., 2021). The Japanese Islands, located east of Eurasia, are composed of the Eurasian, North American, Pacific, and Philippine Sea Plates and were formed approximately 15 million years ago (Barnes 2003). Archaeological evidence suggests that the first human presence in the Japanese Archipelago occurred during the Late Pleistocene, which is 40,000-30,000 years ago. The archipelago experienced two distinct periods in the subsequent Paleolithic and later periods, the Jomon and Yayoi (Hanihara, 1991). These two periods are characterized by lifestyle, cultural, and biological changes associated with prehistoric populations and cultural dispersal events (Hanihara 1991; Hudson 1999; Mizoguchi, 2013; Boer et al., 2020). The Jomon period was characterized by hunter-gatherer-fisher lifestyles, lasting more than 10,000 years, from the final Pleistocene to the Holocene. The transition from the Paleolithic to the Jomon period may have begun with the appearance of pottery use (Taniguchi 2017; Lucquin et al. 2018). However, paddy rice cultivation prospered during the Yayoi period in the Japanese Archipelago, and lifestyle changes are thought to have been introduced by immigrants from the Eurasian continent. Therefore, comparing individual genomes from these two periods provides valuable genetic evidence for considering human diffusion in Eurasia since the Late Pleistocene.

In the genomic analysis of archaeological samples, the quality of genome sequences from ancient remains is subject to environmental conditions, and some data often show a low depth of coverage. Furthermore, the enhancement of ancient human genetic resources is challenging in East Asia, which is warmer and wetter than Europe, northern Eurasia, and North America (Kihana et al., 2013; Kistler et al., 2017; Mizuno et al., 2020). In the Japanese Archipelago, >30-fold high-coverage ancient genomes have been obtained only from Hokkaido (Kanzawa-Kiriyama et al., 2019), which is relatively cold and has a higher latitude than mainland Japan. However, obtaining well-preserved ancient DNA from the mainland of the archipelago, where warm and acidic soils prevail, is challenging. In assessing prehistorical immigrants to Japan, high-coverage ancient genomes on the mainland can elucidate some aspects of the migration history that has occurred in East Asia since the Late Pleistocene.

To enhance the quality of ancient human genetic resources in this region, we obtained high-coverage genomes of individuals from two different periods in the Japanese archipelago. These high-coverage genome sequences allow inferences of several copy number polymorphisms and diploid genotyping and demographic inferences from the individual genome, which have not been reported in previous ancient genomes in this region. Furthermore, our analysis indicated that the Yayoi individual possessed ancestral components found in Eurasian populations, suggesting that genetic lineages exist from the continent that differed from those of the indigenous Jomon people. Our results demonstrate the potential of deep-sequencing approaches for ancient genome research. The analysis of these high-quality ancient genome sequences will shed light on the historical interactions among East Eurasian populations since the Late Pleistocene and the estimation of individual traits from ancient remains.

## 2. Materials and Methods

### 2.1 Archeological sites

#### 2.1.1 Iyai-rock shelter site

The Iyai-rock shelter site is located on the edge of the Jo-shin-etsu mountains between the Shinano and Tone Rivers, which are in the center of Honshu, Japan (Naganohara-cho, Gunma Prefecture; 36°33’28” N latitude, 138°38’50” E longitude) (Figure 1A). The shelter at the site was a rock wall composed of welded tuff. The site has been reported to be an archaeological site dating from the Initial Jomon to the Late Yayoi period, with an open-sloping terrace extending from the interior of the shelter to the front space (Taniguchi and Asakura, 2017). Iyai1 (IY1) was discovered in the summer of 2015, and its nearly complete skeleton was excavated in 2016 (Kondo et al. 2018).

**Figure 1A.**
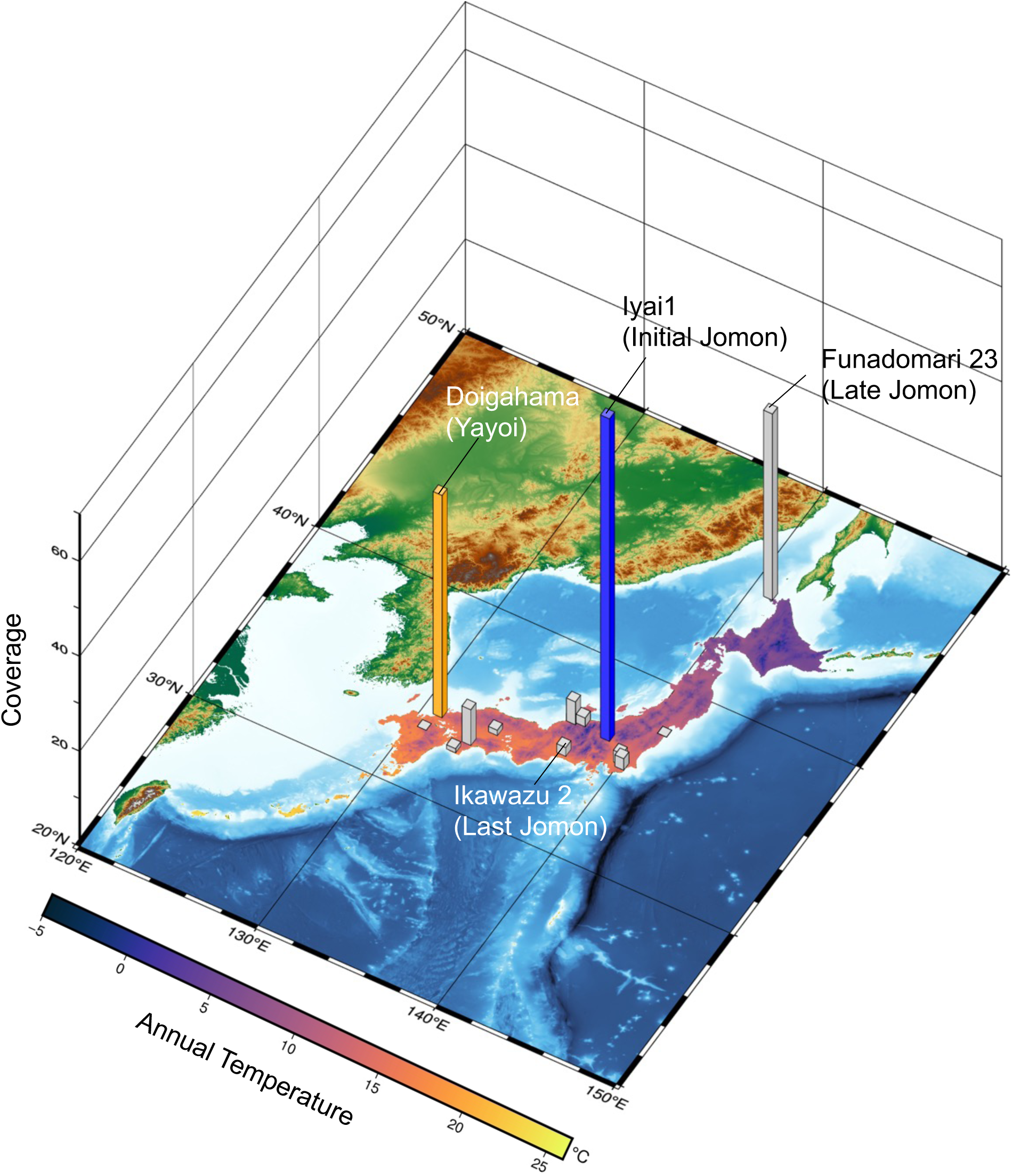
Location of archaeological sites in representative ancient human genomes in the Japanese archipelago. The height of the bar plot indicates the average coverage in the autosomal genome. The colored bars are the high-coverage ancient human genomes (Doigahama and Iyai1) reported in this study. The average temperature is indicated with yellow, higher temperatures, and dark blue, lower temperatures. Annual temperatures are based on mesh-averaged data for 1991-2020 published by the Japan Meteorological Agency.

#### 2.1.2 Doigahama site

The Doigahama site is a cemetery ruin from the early to middle-Yayoi period located on Doigahama beach and is part of southwestern Honshu, Japan (Shimonoseki City, Yamaguchi Prefecture, 34°17’26” N, 130°53’19” E) (Figure 1A). The lime content of shells mixed with sand is suitable for preserving bones, and more than 300 human bones have been found at this site (Igawa et al., 2009).

### 2.2 Experimental samples

We used Jomon individual IY1 from the Iyai-rock shelter site, and Yayoi individual DO from the Doigahama site as samples for whole-genome analysis. Radiocarbon dating indicates that IY1 has a calibrated date of 8,300-8,200 calBP (Kondo et al., 2018), which is the later part of the Initial Jomon period. The DO showed a radiocarbon date of 2,306-2,338 calBP (Mizuno et al., 2021), which belongs to the Middle Yayoi period (Figure 1B). The petrous bones were used for DNA extraction. The bones were sampled using a sterile electric drill cutter (Dremel) and between 130 and 150 mg of bone powder. To prevent contamination with exogenous DNA, DNA extraction, purification, and NGS library preparation were performed in a clean room dedicated to ancient DNA, based on our previous methods (Kihana et al., 2013; Mizuno et al., 2020). NGS libraries were prepared for each extract using single– and double-stranded DNA library preparations for Illumina NGS (Supplementary Table S1). To examine the ancient DNA-like damage patterns of the two samples, we prepared NGS libraries without DNA repair treatments. After confirming postmortem DNA damage patterns, we prepared pretreatment NGS libraries using the PreCR Repair Mix (New England BioLabs) to reduce the impact of postmortem damage on the analysis. These libraries were sequenced using the Illumina HiSeqX system and were used for subsequent genomic analyses (Supplementary Table S1).

**Figure 1B.**
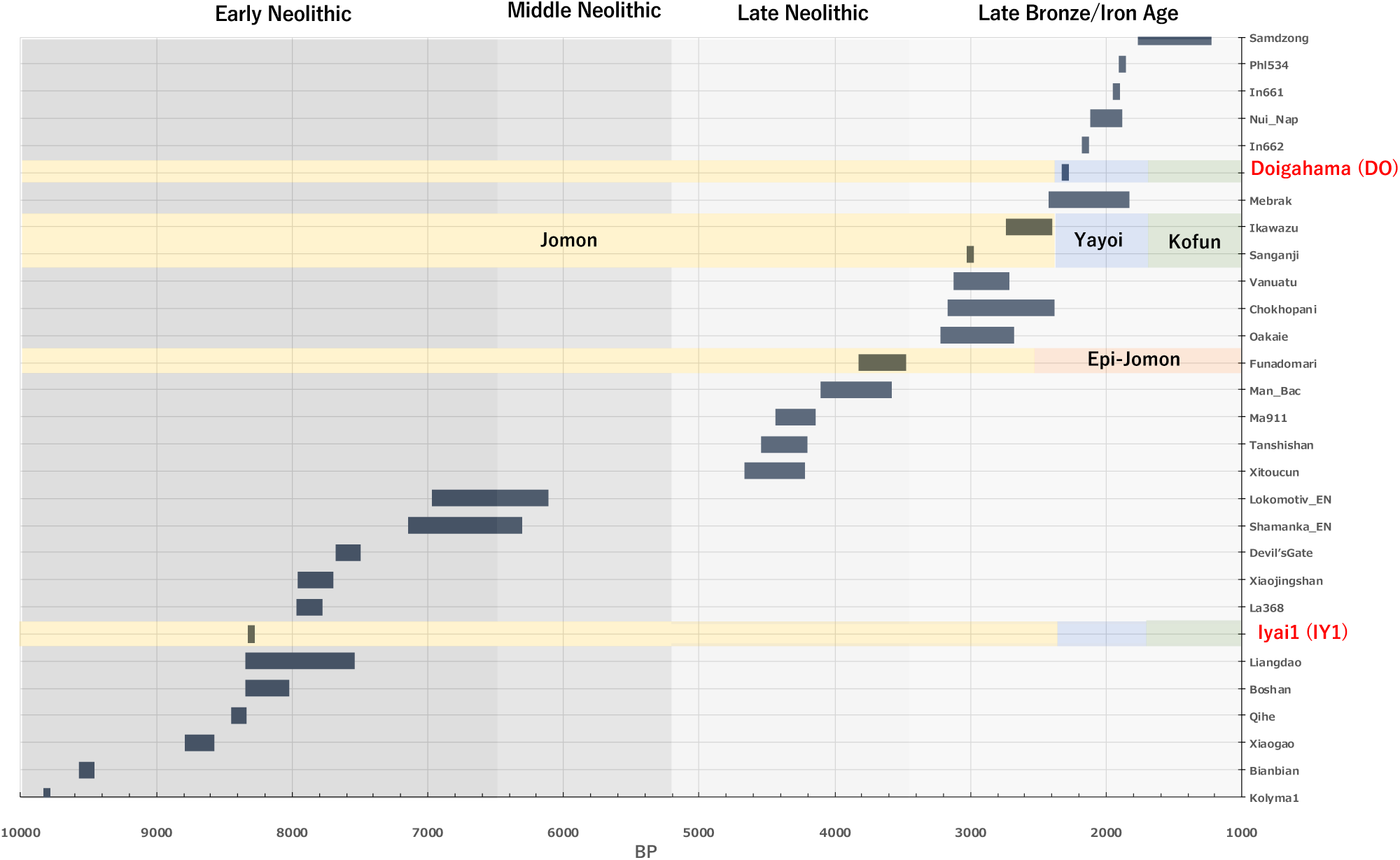
Representative ancient genomes since the Neolithic period in Asia.

**Figure 1C.**
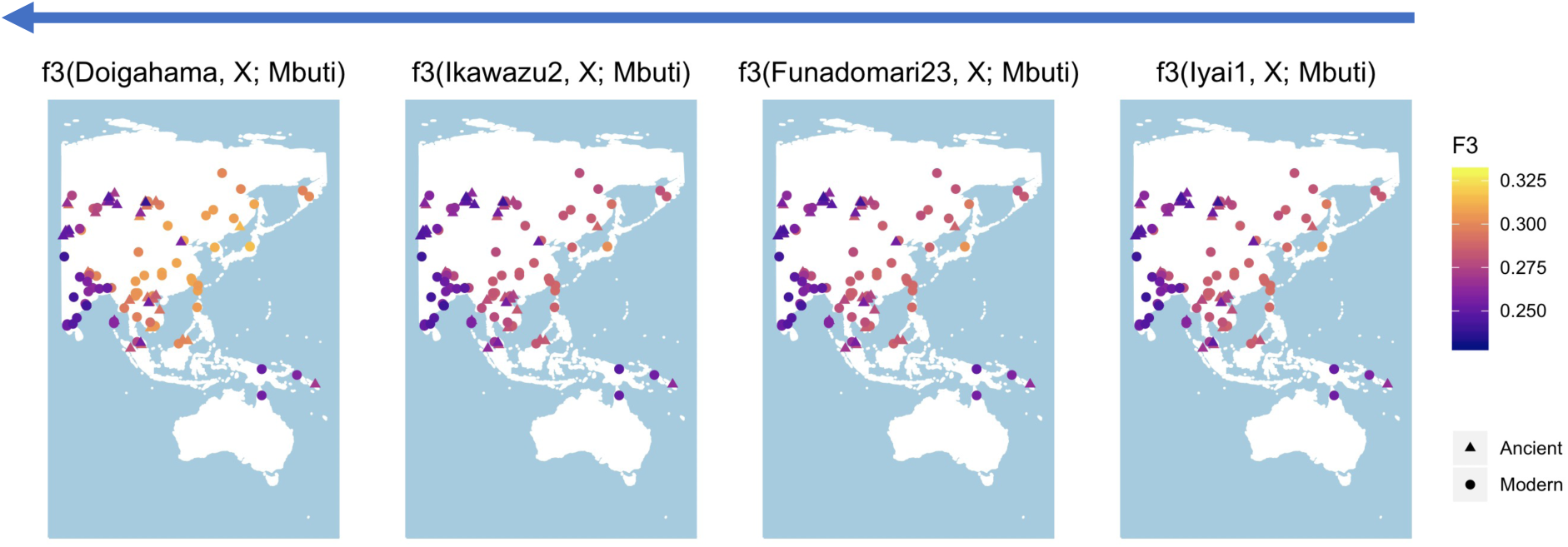
Transformation of genetic affinities among surrounding East-Eurasian and Asian populations. This figure presents a summary of the *f3* values for the representative ancient human genomes (Iyai1, Funadomari23, Ikawazu2, and Doigahama) from the initial Jomon to Yayoi periods in the Japanese archipelago in relation to the surrounding regional populations. The colors correspond to the *f3* values, with brighter ye llow-colored points indicating higher genetic affinity.

### 2.3 Sequence analysis

The raw sequencing reads from each library was processed using FASTP (ver. 0.20.0) (Chen et al., 2018) to remove adapter, low-quality, and read sequences of less than 35 bases. The filtered raw reads were mapped to the human reference genome sequence hg19/GRCh37 using bwa aln (ver. 1.9) (Li et al., 2009) with the parameters for ancient DNA (-l 1024 –n 0.01 –o 2). Duplicate reads were removed from each library using the MarkDuplicates command in the Picard Tool Kit (Broad Institute, 2019). Finally, the alignment reads from each library were merged, and low-quality mapped reads (< MAPQ30) were filtered out using SAMtools (ver. 1.9) (Li et al., 2009b). The average depth of coverage across the genome was determined using the coverage commands in SAMtools.

### 2.4 Biological sex estimation

Ry (Y/X+Y) ratios were calculated based on the number of reads mapped to the human X and Y chromosomes (Skoglund et al., 2013). The biological sex was estimated from the Ry ratios obtained for IY1 and DO.

### 2.5 Maternal and paternal lineage analysis

To assign maternal lineages, the mitochondrial haplogroups of IY1 and DO were estimated using MitoSuite (ver. 1.0.9) (Ishiya and Ueda, 2017). For the male DO sample, the paternal lineage was estimated based on SNPs in the non-recombining region (NRY) of the Y chromosome using Y-LineageTracker (ver. 1.3.0) (Chen et al., 2021). In addition, the NRY SNPs were combined with the worldwide Y-lineage dataset (Hallast et al. 2021), and a Y-chromosome phylogenetic tree was constructed using Y-LineageTracker (ver. 1.3.0) (Chen et al., 2021).

### 2.6 Ancient DNA authenticity

Misincorporation and DNA fragmentation were confirmed using the Mapdamage software (ver. 2.0.8) (Jonsson et al., 2013) with aligned reads from non-repaired NGS libraries. In addition, DNA contamination rates were calculated based on definitive haplogroup sites in the mitochondrial genome using MitoSuite (ver.1.0.9) (Ishiya and Ueda, 2017; Ishiya and Ueda, 2019). In addition, for DO, which is a male individual, autosomal DNA contamination of the nuclear X chromosome was evaluated using ANGSD (ver. 0.939) (Korneliussen et al., 2014).

### 2.7 Variant calling

Single-nucleotide variant calling and haplotype-phase genotyping were performed using GATK (ver. 3.8.1) (McKenna et al., 2010) from reads mapped to the human reference genome sequence. To reduce the impact of false-positive detection, observed mutations in regions masked by the reference sequence, multiallelic sites consisting of three or more alleles, and heterozygous sites with an alternate allele frequency of less than 0.2 were filtered out. Filtered variants were summarized as VCF/PED files and merged into population genotyping data using Plink (ver. 1.90b4) (Purcell et al., 2007). Finally, the coverage of filtered SNPs was calculated using the geno-depth flag in vcftools (ver.0.1.16) (Danecek et al., 2011).

### 2.8 Panel design

The obtained SNPs data were merged against the 1240K/2240 K SNPs global population panel from the Allen Ancient DNA Resource (https://doi.org/10.7910/DVN/FFIDCW; https://doi.org/10.1101/2023.04.06.535797). The merged population panel included 1KGP, HGDP, and the previously reported ancient DNA samples. Related samples were analyzed for kinship using KING (ver. 2.3.2) (Kuhn et al., 2018). Samples that were highly related to each other were excluded from this study. Our population panel consist of 1,444 individuals (Supplementary Table S2), including present-day and ancient samples, located mainly in seven geographical regions (America, Central Asia, East Asia, South Asia, North Asia, Oceania, and the Middle East).

### 2.9 HLA haplotyping

The Human Leukocyte Antigen (HLA) gene complex is one of the most highly polymorphic regions in the human genome. It has a significant impact on the diversity of the immune system. We attempted to determine the HLA typing of Jomon IY1 and Yayoi DO from shotgun sequencing reads. HLA typing was performed based on 4-digit alleles using xHLA (ver.4.0.3) (Xie et al., 2017). The frequencies of the inferred HLA typing in the local population were retrieved from the Allele Frequency Net Database (http://www.allelefrequencies.net/) (Gonzalez-Galarza et al., 2019).

### 2.10 Inference of AMY1 gene copy numbers

A high-starch diet is characteristic of farming and hunter-gathering societies that live in arid environments (Perry et al., 2007). By contrast, hunter-gatherers and pastoralists in colder regions and tropical rainforests consume very low amounts of starch in their diets. The copy number of the amylase gene (AMY1) positively correlates with the level of amylase protein in the saliva. This suggests that these differences in eating styles may exert selective pressure on amylase, an enzyme that hydrolyzes starch. The copy numbers of amylase genes in Yayoi and Jomon individuals were inferred from the depth of coverage of the AMY1 gene region mapped to the human reference sequence using AMYCN (ver. 2020-03-18) (Eisfeldt et al., 2018). The inferred copy numbers were compared with the cumulative distribution of modern hunter-gatherers in cold weather regions, modern Japanese, and pre-agricultural European hunter-gatherers (Labraña 1) (Olalde et al., 2014).

### 2.11 Demographic inference

We used the sequential Markovian coalescence method to infer demographic history from the Yayoi and Jomon individual genomes (Mather et al., 2020). This approach used ancient Japanese samples called diploid sequences from the mainland Jomon individual (IY1) and the Yayoi individual (DO). Whole-genome diploid consensus sequences were obtained from aligned reads using bcftools (ver.1.9) (Li, 2011), and these sequences were used as input values to infer demographic history. Pairwise sequential Markovian Coalescent inference was conducted using PSMC (ver. 0.6.5-r67) (Li and Durbin, 2011). More recent demographic inferences using unphased genomes were performed using the SMC++ program (ver. 1.15.2) (Terhorst et al., 2017).

### 2.12 LD pruning

To extract SNP markers that were not in linkage disequilibrium, we performed LD pruning using plink (ver. 1.90) (Purcell et al. 2007) with the option “--indep-pairwise 50 10 0.1.”

### 2.13 Principal component analysis

To investigate the genetic relationships of Jomon (IY1) and Yayoi (DO) individuals to worldwide populations, principal components analysis (PCA) was performed using the smartpca program in the EIGENSOFT package (ver.7.2.1) (Patterson et al., 2006). The analysis was carried out with the outlier removal option off (set “numoutlieriter” as zero), normalization options, and the lsqproject on to ensure that no samples were automatically excluded.

### 2.14 Unsupervised clustering analysis for ancestral genetic structure

We used Admixture (ver.1.3.0) (Alexander et al., 2009) to perform unsupervised genetic clustering of modern worldwide populations with ancient samples from the Japanese Archipelago. In addition, to compare ancestral proportions among samples from regional populations, we performed an admixture analysis using samples clustered around East Asia, including the Jomon and Yayoi individuals in PCA. *K*=2–12 clusters for worldwide population panels and *K*=2–12 clusters for regional populations were explored using ten independent trials with 5-fold cross-validation. We repeated the admixture analysis ten times for each *K* value to verify that the estimation converged to the same state. The repeated results were summarized using CLUMPAK (ver. 1.1.2) (Kopelman et al. 2015).

### 2.15 *f3*-statistics

To assess genetic affinity among populations, outgroup-*f3* statistics were performed using the qp3pop (ver. 980) program of AdmixTools (Patterson et al., 2012). The outgroup-*f3* measures the shared genetic drift between the source populations, X and Y, compared with the outgroup population (Mbuti). To exclude the effects of ancient DNA damage, we generated genome-wide single nucleotide polymorphism (SNPs) data restricted to transversion sites and performed a similar analysis.

### 2.16 Inference of population divergences

To infer the maximum likelihood trees using the proposed admixture models, TreeMix (ver. 1.13) (Pickrell et al., 2012) with a population subset of ancient and present-day individuals across East Eurasia and Asia: Ust_Ishim, Sunghir_UP, Yana_UP, MA1, Tianyuan, Papuan, Onge, USR, Jomon, Shamanka_EN, Chokhopani, Ami, CHB, Yayoi (DO), JPT (1 KGP +HGDP), Devil’s Cave_N, and Kolyma. Mbuti was the outgroup at the root of the tree, and the migration event (*m*) was verified to be *m*=0 to 5. Single nucleotide polymorphisms (SNPs) at the transition sites were used to estimate the maximum likelihood of phylogenetic tree construction and migration event estimation to exclude the effects of ancient DNA misincorporation.

### 2.17 Inference of admixture timing

Although the actual demographic process of the admixtures is complex, simple models can provide valuable insights into their characteristics. Here, we used ALDER (ver.1.0.3) (Loh et al., 2013), based on admixture-induced linkage disequilibrium (LD), to infer the admixture timing between individuals and populations by comparing weighted LD curves using reference populations. The target population was modern Japanese, and modeling was performed using IY1 as the resource population representing Jomon ancestry, Yayoi individual DO, and modern humans from mainland China (Han) as immigrant/continental population sources.

### 2.18 Admixture modeling

Admixture modeling was performed using the qpGraph program in Admixtools (ver. 7.0.2) (Patterson et al. 2012; Haak et al. 2015) to infer plausible admixture models between populations of interest in the East Asian lineages. Modeling was performed using the following samples: Dinka, Malta, Onge, MA1, Shamanka, Alaska_LP, USA_Ancient_Beringian, Chokhopani, Japan_Jomon_Iyai, and Japan_Yayoi_Doigahama. In addition, we used the qpAdm program in Admixtools (ver.7.0.2) (Haak et al., 2015), to assess immigrant sources that contributed to modern Japanese. Admixture proportions were estimated in the model when one source population of the modern Japanese population was fixed to the Jomon and the other to ancient East Asian and Far Eastern populations, including the Yayoi individual. We used genome-wide transversion sites for this analysis and ran with the option “allsnps: YES”. The following nine populations were used as outgroups: Mbuti, Russia_Ust_Ishim, China_Tianyuan, Russia_Kostenki14, Papuan, ONG, Yana_UP, Liangdao2, and Yumin. To detect independent gene pools, the independence in these outgroup and reference populations was assessed using the qpWave program in Admix tools (ver. 7.0.2) (Patterson et al. 2012; Haak et al. 2015).

### 2.19 Inference of introgressed fragments from archaic hominins

To infer ancestry fragments derived from archaic hominins (Neanderthal or Denisovan) across each genomic position in Jomon and Yayoi individuals, we attempted to fit the allele frequency by comparing it with reference sources based on a Hidden Markov Model (Peter, 2020) (https://github.com/BenjaminPeter/admixfrog). The reference resources used here are Neanderthals and Denisova from Vindija and Altai and Yoruba from the African population. We set the “--c0” option because we assumed low-level contamination in our dataset.

## 3. Results

### 3.1 High-coverage ancient genomes from prehistoric Japan

In the Japanese Archipelago, high-quality ancient genomes were obtained from Hokkaido (Kanzawa-Kiriyama et al., 2019), which is relatively cold and has a higher latitude than mainland Japan. However, a high-coverage ancient genome has not yet been reported because obtaining well-preserved DNA remains from the archipelago, where warm and acidic soils prevail, is challenging. In ancient samples, excluding those from Hokkaido in the Japanese Archipelago, the coverage of SNPs and autosomes has often been reported to be low. For example, in the mainland (Honshu), the genome coverage of Ikawazu Jomon (IK002) (McColl et al., 2018; Gakuhari et al., 2020) was 1.85-fold, while in Rokutsu (Wang et al., 2021), and Sanganji Jomon (Kanzawa-Kiriyama et al., 2016) individuals had coverage range of 0.16-2.42-fold, and 0.009-0.014-fold, respectively (Supplementary Table S3). In Kyushu, a southern Japanese island, the autosomal coverage of the Kuma-Nishioda Yayoi individual (YAK002) (Robeets et al., 2021) was also limited to <0.0006-fold (Supplementary Table S3). These low-coverage genomes limit analytical methods and statistical evaluations. Furthermore, the origin of the Yayoi period should be discussed more carefully. For example, regional differences and variations were observed in facial traits in the Yayoi period, as well as human skeletons in the Kyushu region, and the possibility of admixture with indigenous lineages must be considered in this resource. For this origin inference, working with samples from the initial Jomon period, which are closer to the basal part of the Jomon culture, and from the “migratory” Yayoi period, is crucial.

In this study, we performed high-coverage genome sequencing of specimens from the initial Jomon period (IY1) and the middle Yayoi period (DO) in central and southwestern mainland Japan (Figure 1A). IY1 is one of the oldest buried human remains in the Initial Jomon period in the Japanese archipelago. The Doigahama site, from which the DO individual was excavated, is also known as a site for migratory Yayoi people, and the individual could be a representative genomic resource for migratory Yayoi people. Genomic DNA was extracted from the petrous bones of both samples, and the shotgun-sequenced genomes showed high coverage (IY1: autosome-avg. 67.94-fold; DO: autosome-avg. 46.82-fold) (Table 1, Supplementary Figure S1-S3, and Supplementary Table S3). Biological sex was estimated from the read ratio (Ry) mapping to sex chromosomes as female for IY1 (Ry=0.001) and male for DO (Ry=0.090) (Table 1). To validate ancient DNA authenticity, obtained sequencing reads were confirmed to show postmortem deamination patterns (Supplementary Figure S4), and further, both also showed <1% exogenous contaminated DNA (IY1: 0.727%, DO: 0.234% in the human mitochondrial genome; the male individual DO: 0.420% in X-chromosome genome). Maternal mitochondrial haplogroups of IY1 and DO were N9b and D4b2b1, respectively (Table 1). The results of the mitochondrial haplogroup assignments were consistent with those of previous mitochondrial genome studies (Mizuno et al., 2020; Mizuno et al., 2021). The Y chromosome haplogroup of male DO individuals was assigned to the D1a2a lineage, observed in modern Japanese individuals (Supplementary Figure S5 and S6). These two individuals were integrated into a published genome dataset of modern and ancient individuals from this area to delve into the genetic context and ancestral links among the Eurasian people. Genetic kinship analysis confirmed that the two ancient individuals were unrelated to any individuals in the dataset (Supplementary Figure S7).

**Table 1.**
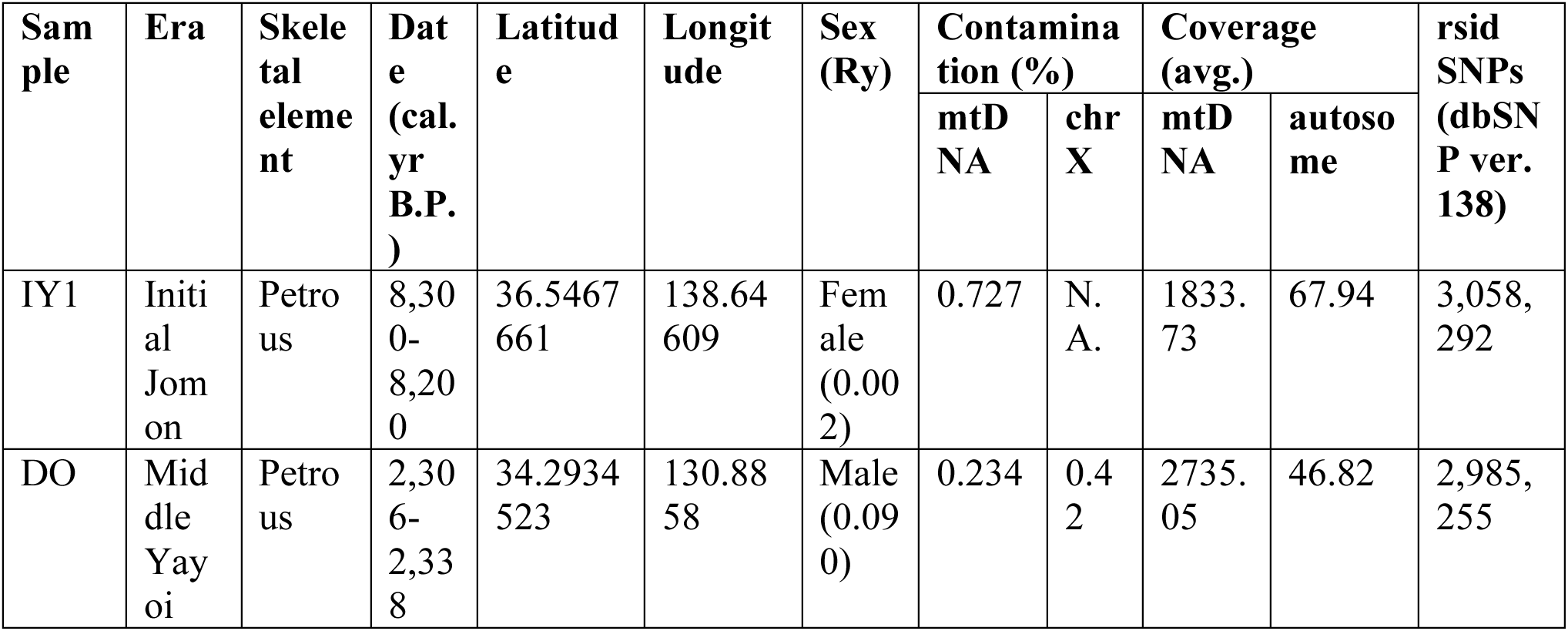
Summary of sequenced samples in this study.

### 3.2 Genetic relationships with worldwide populations

To make inferences about genetic relationships with diverse populations from geographically distant parts of the world, genome-wide SNPs in IY1 and DO were integrated into a worldwide population dataset. Principal component analysis (PCA) was performed using these integrated panels to examine the genetic context of the Jomon and Yayoi individuals (Figure 2). The results (PC1 and PC2) showed that the two individuals (IY1 and DO) were located close to the present-day East Asian population, but the two principal components differentially characterized IY1 and DO. In PC3 and PC4, we were able to distinguish Jomon from other Eurasian populations. IY1, an initial Jomon individual from the mainland, was located near previously reported Jomon individuals close to the Late Jomon individuals from the mainland (ca. 1500-1000 BC). However, IK002 (2720–2418 cal BP) from the Late-final Jomon Period on the mainland and F23 from the Late Jomon Period in Hokkaido (4025 ± 20 (BP ± 1σ)) were located slightly farther away. The Yokchido individual from ancient Korea was included in the Jomon cluster. The Yayoi individual, DO, were closer to the East Asian population than the Jomon individuals.

**Figure 2A.**
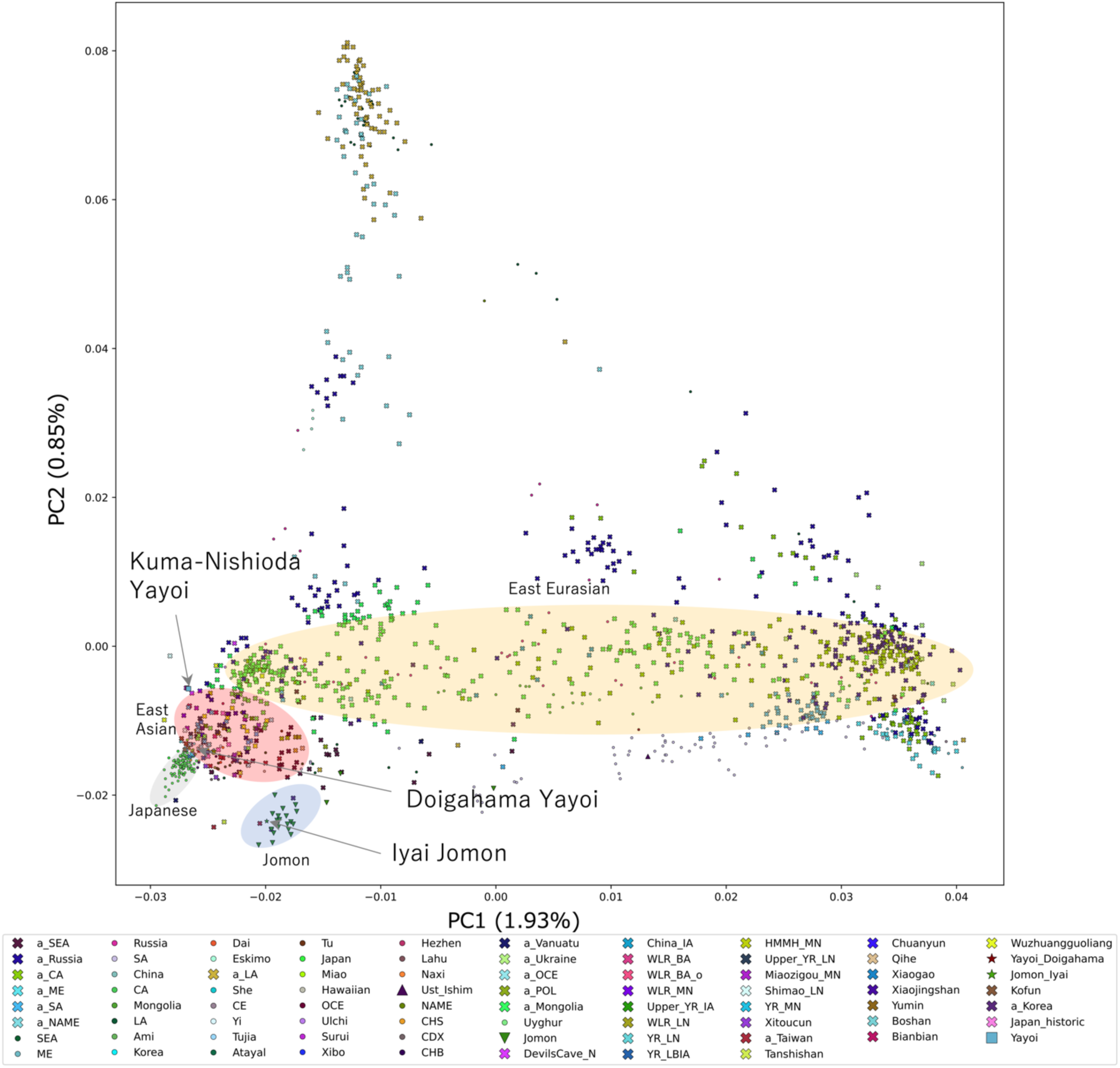
Principal component analysis (PC1 and PC2). This figure shows the results of a principal component analysis using genome-wide SNPs at transversion sites. The values of PC1 and PC2 map each population in the principal component analysis. The colors and shapes of the figure correspond to different populations. The percentage on each axis is the explained variance, which indicates how much of the total variance in the data set is captured by the principal components.

**Figure 2B.**
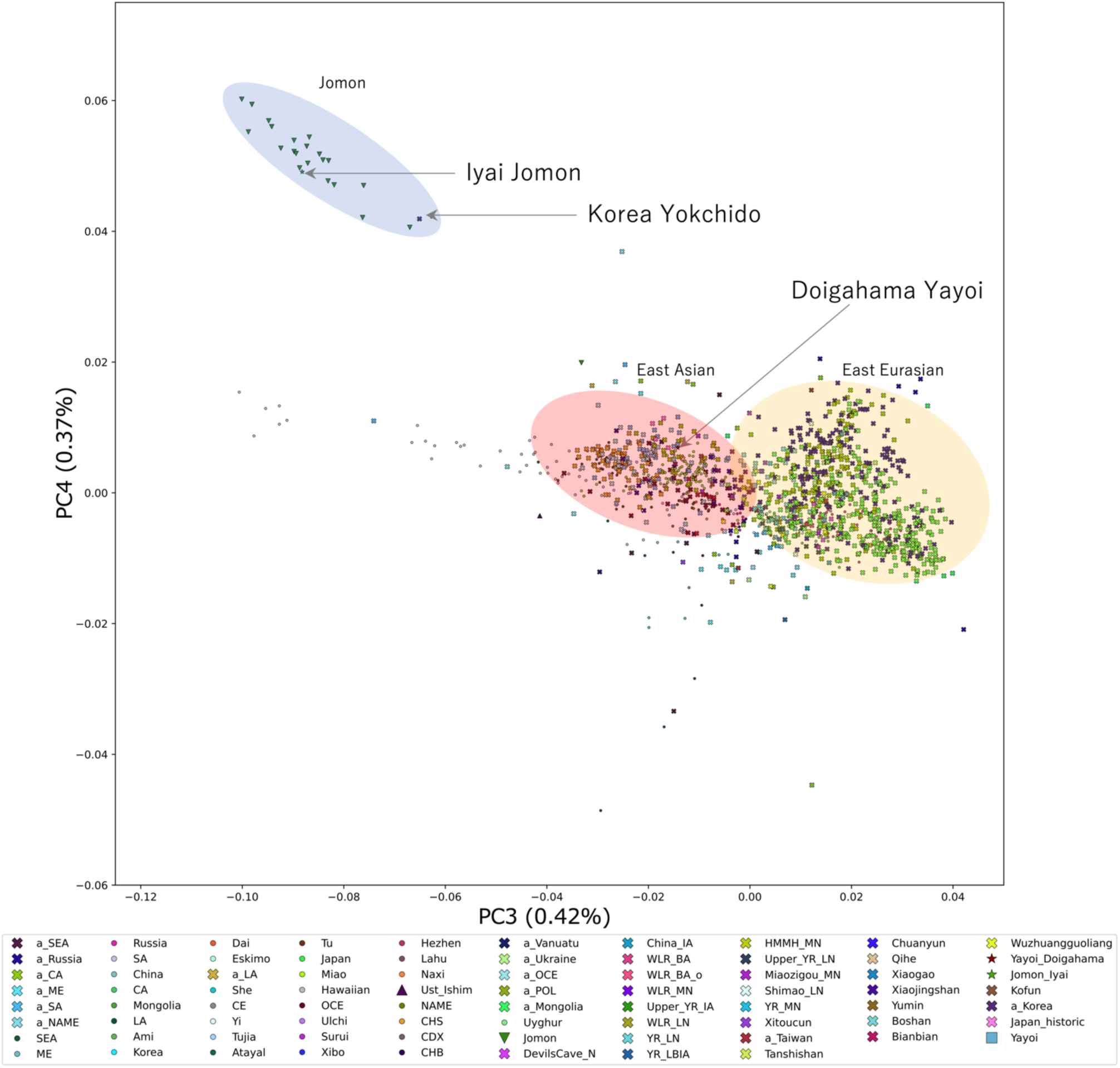
Principal component analysis (PC3 and PC4). This figure shows the results of a principal component analysis using genome-wide SNPs at transversion sites. The values of PC3 and PC4 map each population in the principal component analysis. The colors and shapes of the figure correspond to different populations. The percentage on each axis is the explained variance, which indicates how much of the total variance in the data set is captured by the principal components.

### 3.3 Ancestral contexts of Jomon and Yayoi individuals

To infer the ancestral components of the Jomon and Yayoi individuals, model-based unsupervised clustering analysis was performed using ADMIXTURE (Figure 3). We found that the Jomon and Yayoi individuals shared the same components found in the present-day Japanese population yet were also shown to have several different components (Figure 4). The blue component (P1) observed in Yayoi was not observed in the Jomon individuals. Nonetheless, it was observed at a high frequency in ancient Northeast Asia (ANEA), mainly in Mongolia and the Baikal region. A green component (P4) was also observed in the high-coverage Mainland Jomon IY1 and Hokkaido Jomon (F23). In contrast, this component was not observed in other Jomon individuals (Figure 3). These P1 and P4 components have also been observed in present-day Koreans. This green component is also widely observed in southern China, Southeast Asia, and Oceania populations. Interestingly, we also found that the green components were present at high frequencies in Papuans and Vanuatus individuals (Figure 4; Supplementary Figure S9). The light blue component (P5) was the major component in Jomon; P5 was also observed outside prehistoric Japan in Yokchido individual from ancient Korea (Figure 3).

**Figure 3.**
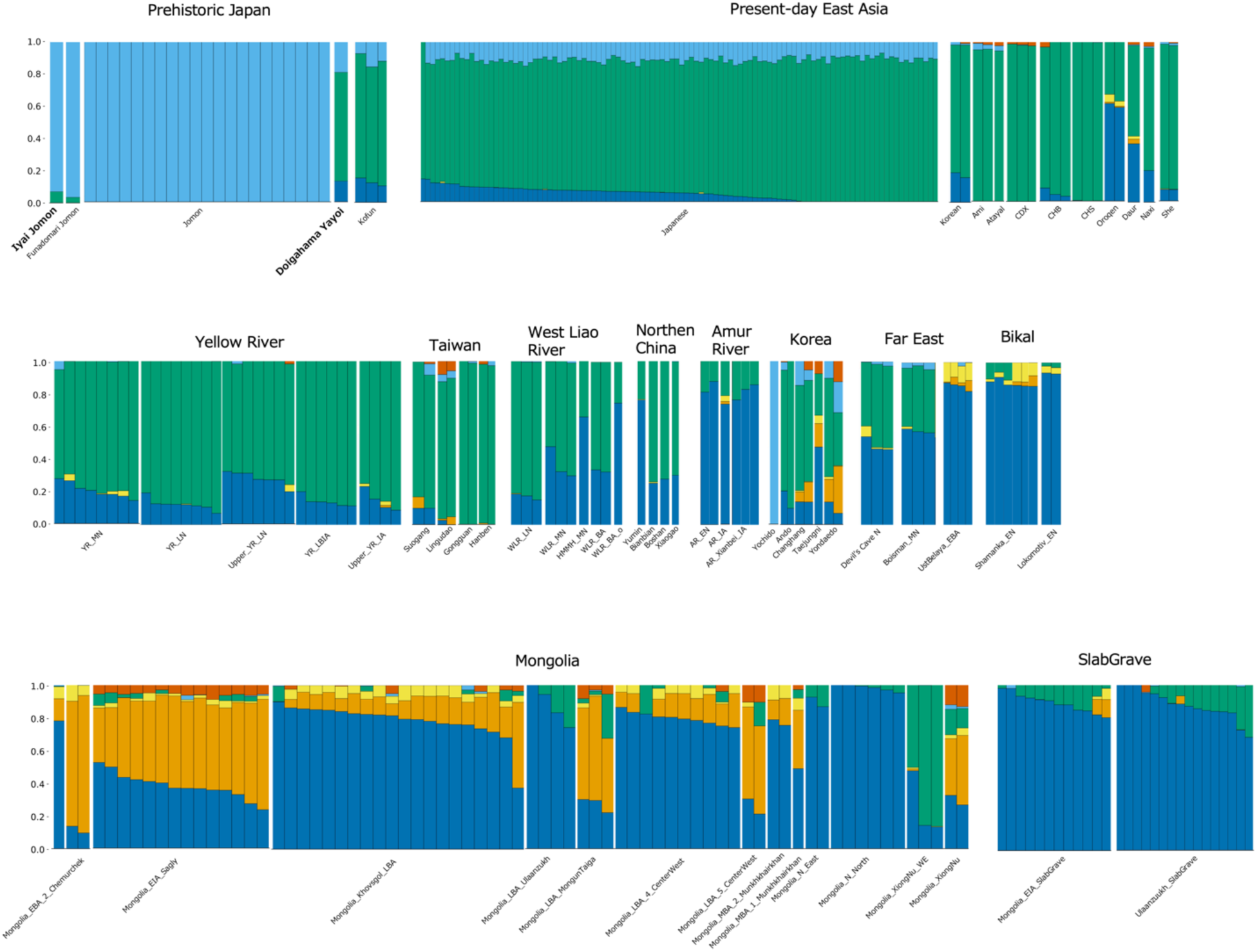
The bar plots of selected individuals from an admixture analysis. The number of ancestor components was validated from *K*=2 to 12, and *K*=6 showed the lowest cross-validation error (Supplementary Figure S8). The ancestor components correspond to the colors P1 (dark blue), P2 (orange), P3 (yellow), P4 (green), P5 (light blue), and P6 (red), respectively.

**Figure 4.**
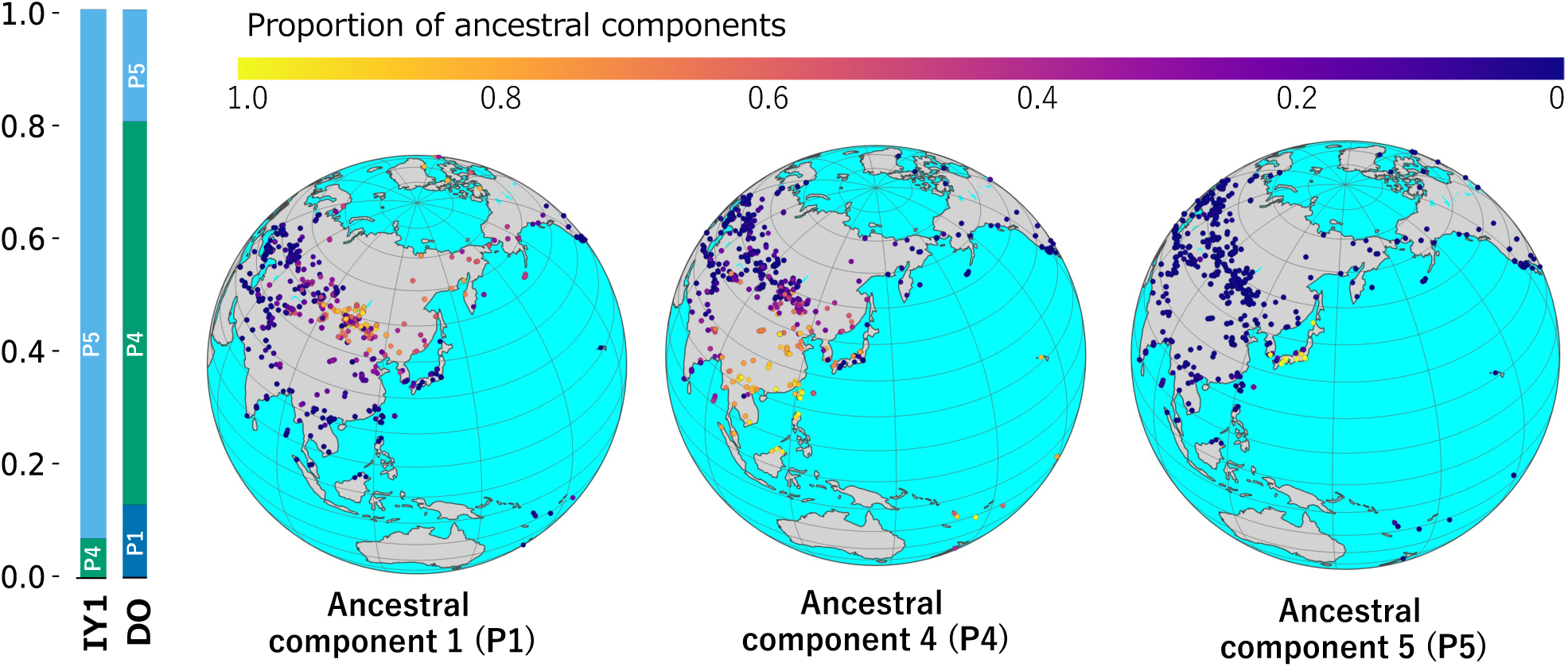
Geographic distribution of ancestral components observed in IY1 and DO individuals. This figure summarizes the bar plots and geographic distribution of the ancestral components observed in the Jomon (IY1) and Yayoi (DO) individuals in the Admixture analysis at *K*=6. The observed ancestral components are marked with P1, P4, and P5, respectively. The colored bars at the top indicate the proportion of each ancestral component observed in each geographic population. The blue component (P1) observed in the Yayoi was not observed in the Jomon individual, but was observed at high frequency in ancient Northeast Asia (ANEA), mainly in Mongolia and the Baikal region. A green component (P4) was also observed in the high-coverage Mainland Jomon IY1 and Hokkaido Jomon (Funadomari 23). This green component is widely observed in southern China, Southeast Asia, and Oceania populations. The light blue component (P5) is the major component in the Jomon; P5 is also observed outside prehistoric Japan in Yokchido individual from ancient Korea.

The outgroup-*f3* and *f4* statistics were used to examine whether these two individuals showed genetic affinities for Eurasian and East Asian populations (Figure 1C, 5, and 6). The genetic affinities against the other Eurasian populations did not change significantly from the early to Late Jomon period; however, itshowed an alteration from the Yayoi period (Figure 1C). To investigate the impact of prehistoric migration events that have been assumed to have occurred in the Japanese Archipelago since the Yayoi period, we focused on *f3* and *f4* values for the surrounding population, especially the Eurasian and Asian populations. IY1, an individual from the initial Jomon period, showed the highest affinity for other individuals belonging to the Jomon period compared with other regional modern populations and ancient Asian individuals (Figure 5). In the *F3* values, the DO of the middle Yayoi individual showed a pattern distinctly different from that of the Jomon individual. The DO showed a higher affinity for Neolithic Far East individuals, such as the Boisman and Devil’s Gate, than for ancient northern/southern Chinese and present-day East Asians (Figure 5 and 6). Furthermore, the Yayoi individual showed affinities with samples from the Iron Age Yankovsky culture (Figure 5). However, the *f4* statistic using Kofun samples, which were considered a migratory resource in a previous study (Cooke et al., 2021), showed slightly different associations with ancient Eurasian individuals in *f4* values (Z>3) (Supplementary Figure S11). We also compared Yayoi and Kofun to their neighboring ancient Eurasian populations and their genetic affinities to ancient human individuals using the outgroup *f3* statistic. The Yayoi individual showed high affinities to Jomon individuals and those of Neolithic and Middle Neolithic Far Eastern ancient samples (Figure 6).

**Figure 5.**
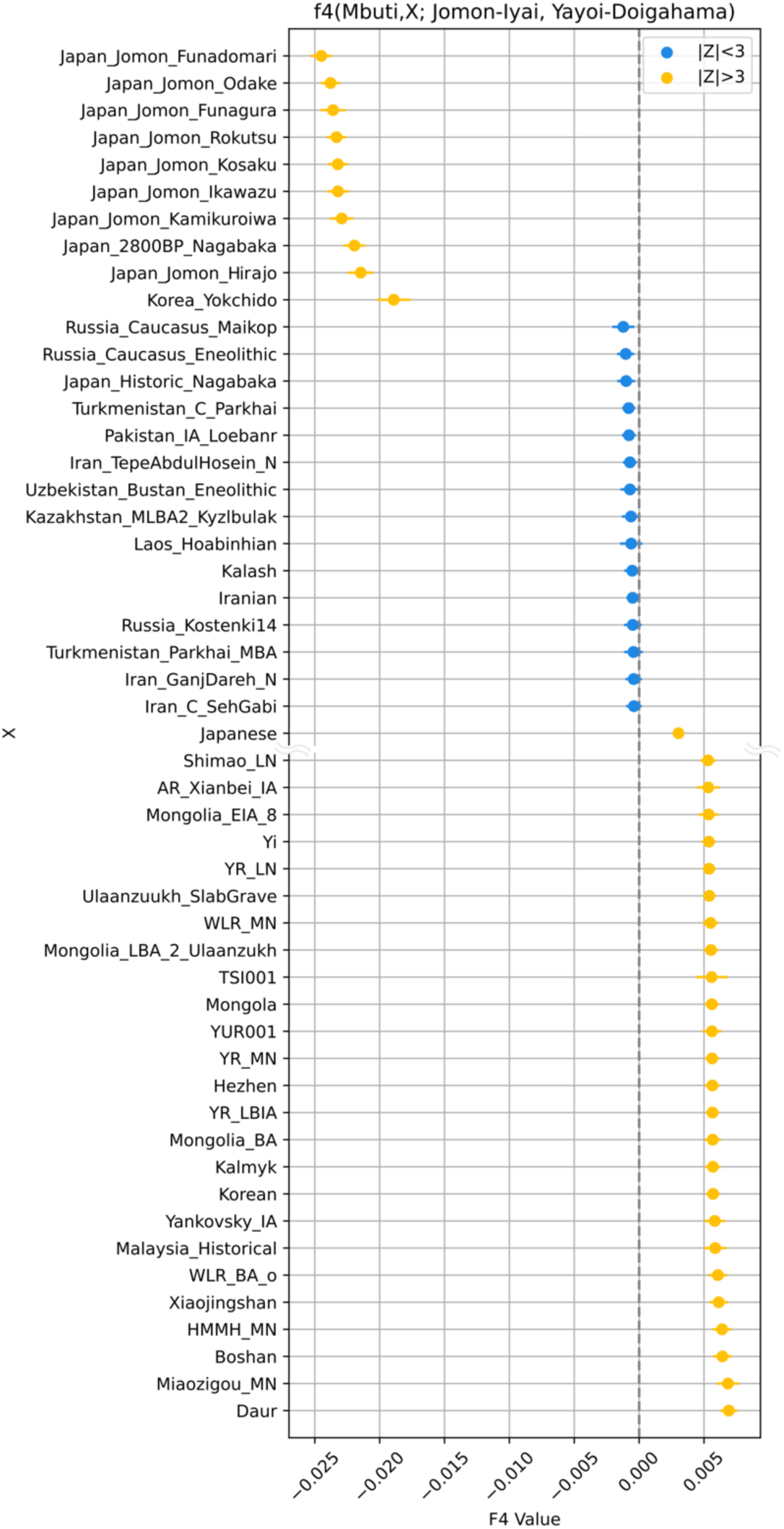
Results of *f4*-statistics highlighting the shared genetic drift between Initial-Jomon (Jomon-Iyai) and Middle-Yayoi (Yayoi-Doighama). In *f4*(Mbuti; X, Jomon-Iyai, Yayoi-Doighama), the top 50 individuals/populations were selected from those with the highest absolute *f4* values. Error bars indicate the standard deviation; absolute values of *Z* greater than 3 are shown in yellow, and values less than 3 are in blue. Negative values indicate more shared genetic drift with Jomon-Iyai and positive values with Yayoi-Doighama.

**Figure 6.**
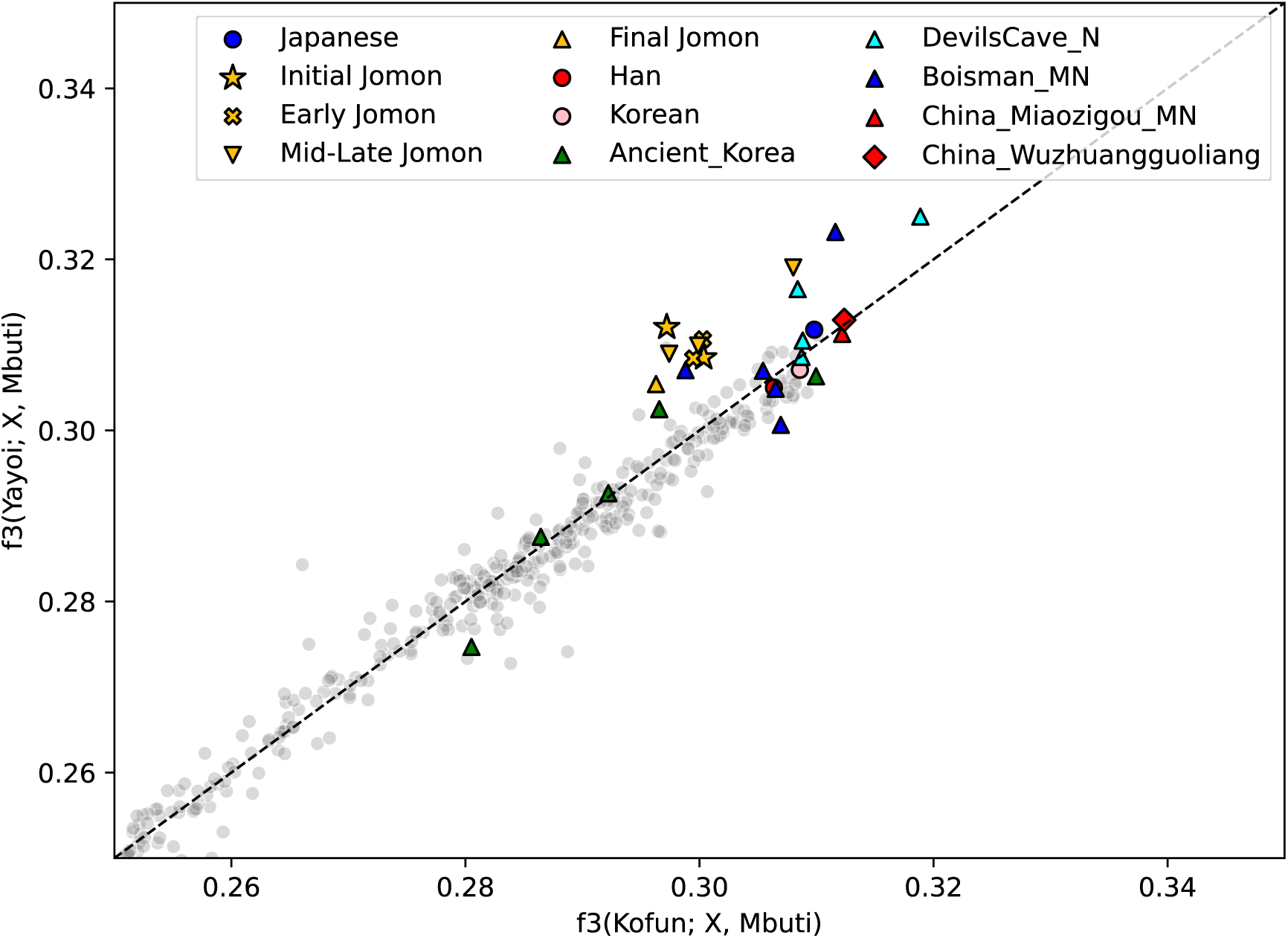
A comparison of outgroup-*f3* statistics with the Yayoi and Kofun individuals against East Asians.

### 3.4 Ancestral contexts of Jomon and Yayoi individuals

To infer the ancestral components of the Jomon and Yayoi individuals, model-based unsupervised clustering analysis was performed using ADMIXTURE (Figure 3). We found that the Jomon and Yayoi individuals shared the same components found in the present-day Japanese population but were also shown to have several different components (Figure 4). The blue component (P1) observed in the Yayoi was not observed in the Jomon individuals, but was observed at high frequency in Ancient Northeast Asia (ANEA), including Mongolia and the Baikal region. A green component (P4) was also observed in the high-coverage Mainland Jomon IY1 and Hokkaido Jomon (F23), respectively. In contrast, this component was not observed in other Jomon individuals with low-coverage genomes (Figure 3). These P1 and P4 components have also been observed in present-day Koreans. The green component (P4) also is widely observed in populations from southern China, Southeast Asia, and Oceania. Interestingly, we also found that the green components are high frequency in Papuans and Vanuatu individuals (Figure4; Supplementary Figure S9). The light blue component (P5) is observed as the major component in the Jomon; P5 is also observed outside the prehistoric Japan in Yokchido individual from ancient Korea (Figure 3).

The outgroup-*f3* and *f4* statistics were used to examine whether these two individuals showed genetic affinities to Eurasian and East Asian populations (Figure 1C, 5 and 6). The genetic affinities against the other Eurasian populations did not change remarkably from the initial to Late-final Jomon period, but clearly showed an alteration from the Yayoi period (Figure 1C). To investigate the impact of prehistoric migration events that have been assumed to have occurred in the Japanese archipelago since the Yayoi period, we focused on *f3* and *f4* values with the surrounding population, especially Eurasian and Asian populations. *F3* values with IY1 are high among other Jomon individuals and show no significant affinities with other continental individuals (Figure5). In contrast, the middle Yayoi individual DO showed a distinctly different pattern from the Jomon individuals. The DO showed a higher affinity with Neolithic Far East individuals such as Boisman and Devil’s Gate than ancient northern/southern Chinese and present-day East Asians (Figure5 and 6). Furthermore, the Yayoi individual also show affinities with samples from the Iron Age Yankovsky culture (Figure5). However, in the *f4* statistic using Kofun samples, which was considered one of the migratory resources in the previous study (Cooke et al., 2021), showed slightly different associations with ancient Eurasian individuals in *f4* values (with Z>3) (Supplementary Figure S11). We also compared Yayoi and Kofun to their neighboring ancient Eurasian populations or their genetic affinities to ancient human individuals, respectively, using the outgroup *f3* statistic. The Yayoi individual showed affinities to Jomon individuals, as well as those of Neolithic and Middle Neolithic Far East ancient samples (Figure 6).

### 3.5 Ancestral demographic changes

To estimate the population size transitions that occurred in the past, we performed demographic inferences based on the Coalescent Theory. Our results highlight the similarities and differences in population sizes in the Jomon and Yayoi populations during their ancestral periods. The population sizes of the two groups were not discernibly different until approximately 100,000 years ago, after which the Jomon ancestral population experienced a notable increase until roughly 20,000 years ago (Figure 7A). These differences between the two groups were more pronounced after the last glacial maximum (LGM). The estimates from the Jomon individual showed a marked decline in population size after the LGM. In contrast, the Yayoi individual showed a gradual increase in population size after the last glacial period (Figure 7B).

**Figure 7A.**
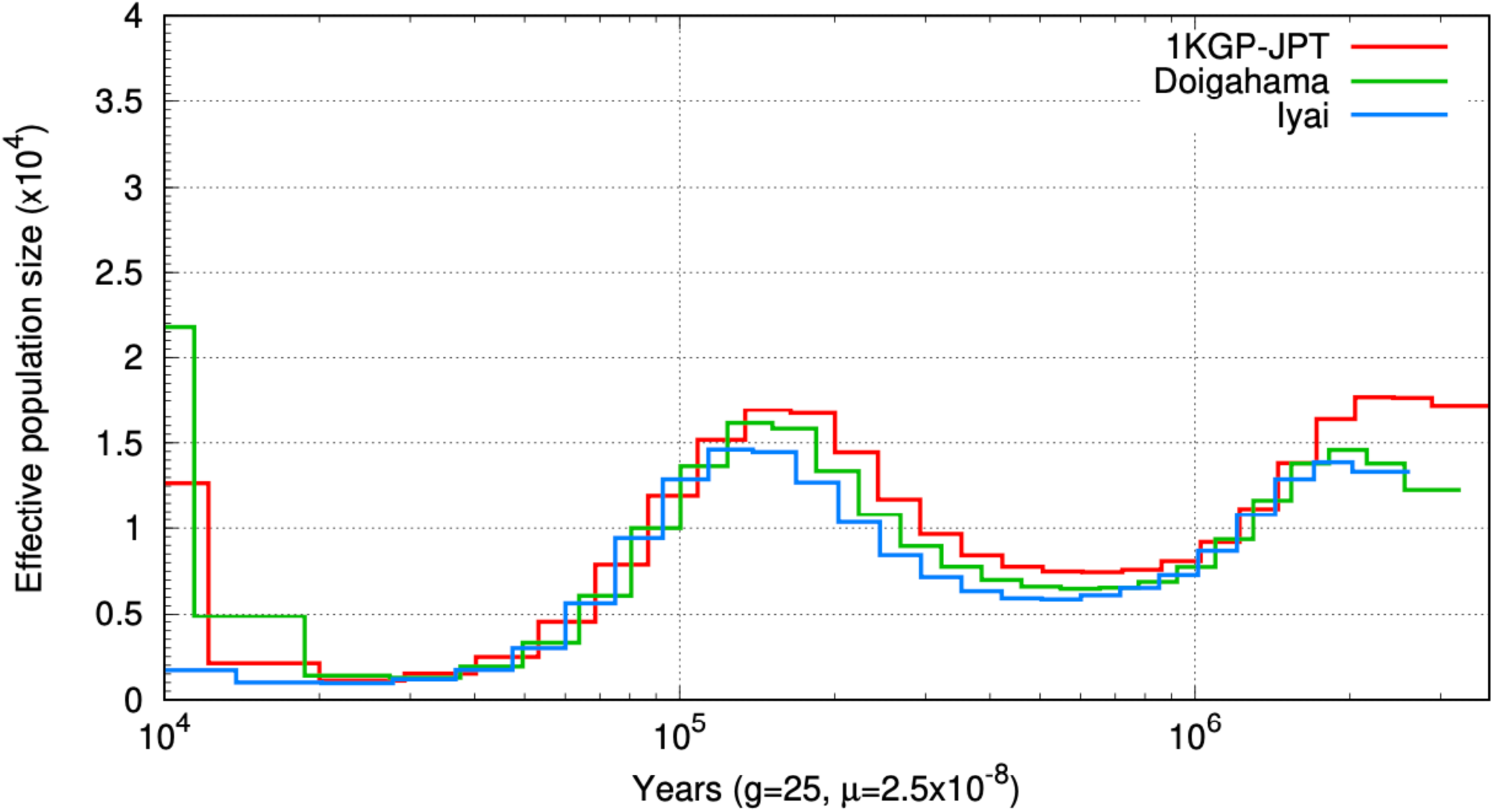
The past demographic inferences for ancient and modern Japanese individuals. This figure shows past demographic plots inferred from the genome of a single individual. The red line represents 1KGP-JPT (modern Japanese), and the green and blue lines represent Yayoi and initial Jomon individuals, respectively. Here, we assume 25 years per generation and a mutation rate *μ* of 2.5 × 10^-8^.

**Figure 7B.**
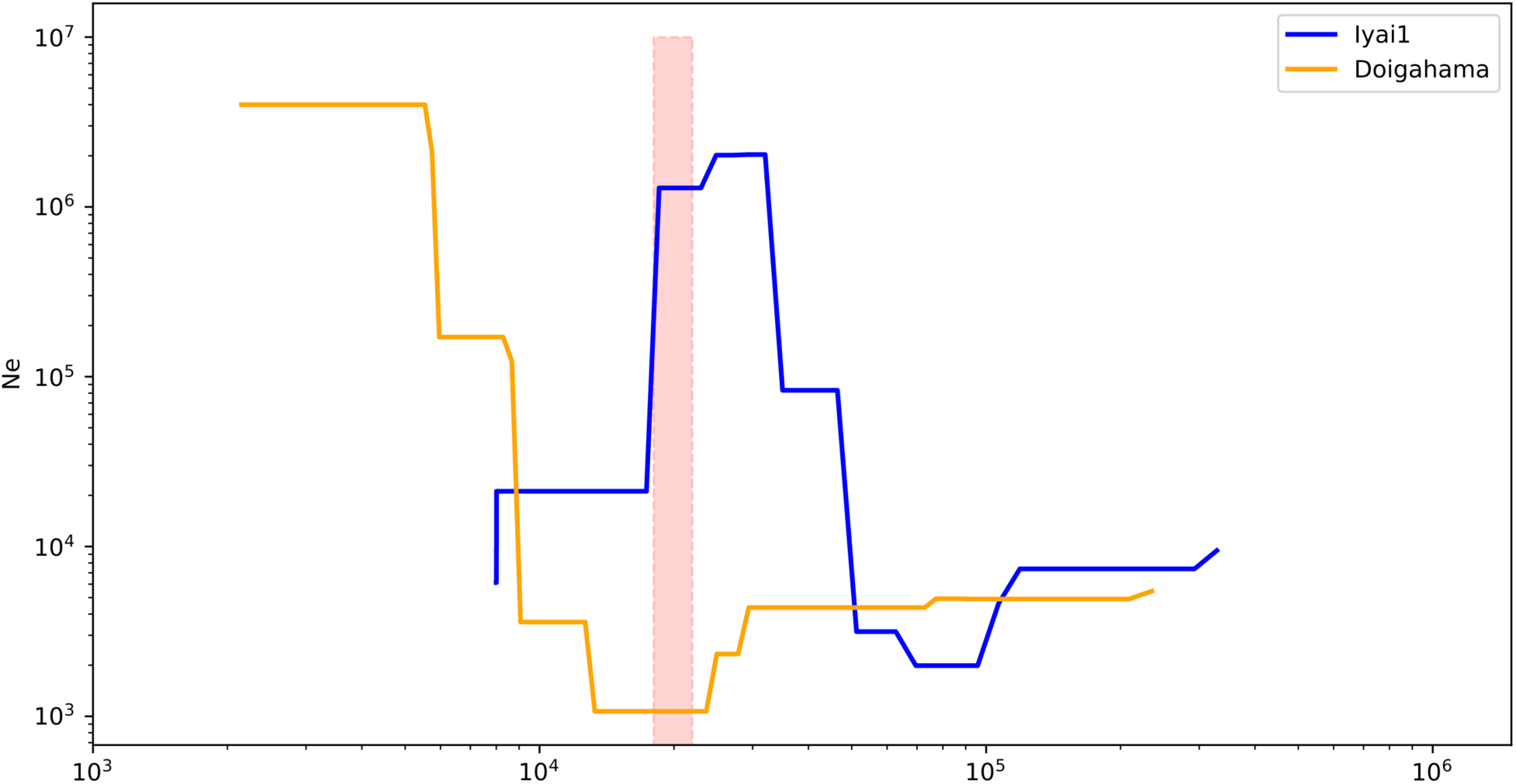
The more recent past demographic inference in ancient Japanese individuals. This figure shows recent demographic shifts, including after 10,000 years, estimated from the genome of a single individual. The pink shade indicates the last glacial maximum (LGM). The blue and yellow lines indicate the initial Jomon and middle-Yayoi individuals, IY1 and DO, respectively.

We further inferred runs of homozygosity (ROH) for several individuals ranging from Jomon to Kofun to detect differences in population size (Figure 8). Some Jomon individuals showed a gradual decrease in ROH from the initial to middle Jomon. In contrast, several individuals with high ROH were found in the Late to Late-final Jomon period. This change may be due to regional characteristics and environmental or climatic changes like cold weather conditions. The estimated ROHs of the Yayoi and Kofun samples were lower than those of the Late and Late-final Jomon individuals. Continental migration events may have contributed to the decreased trends.

**Figure 8.**
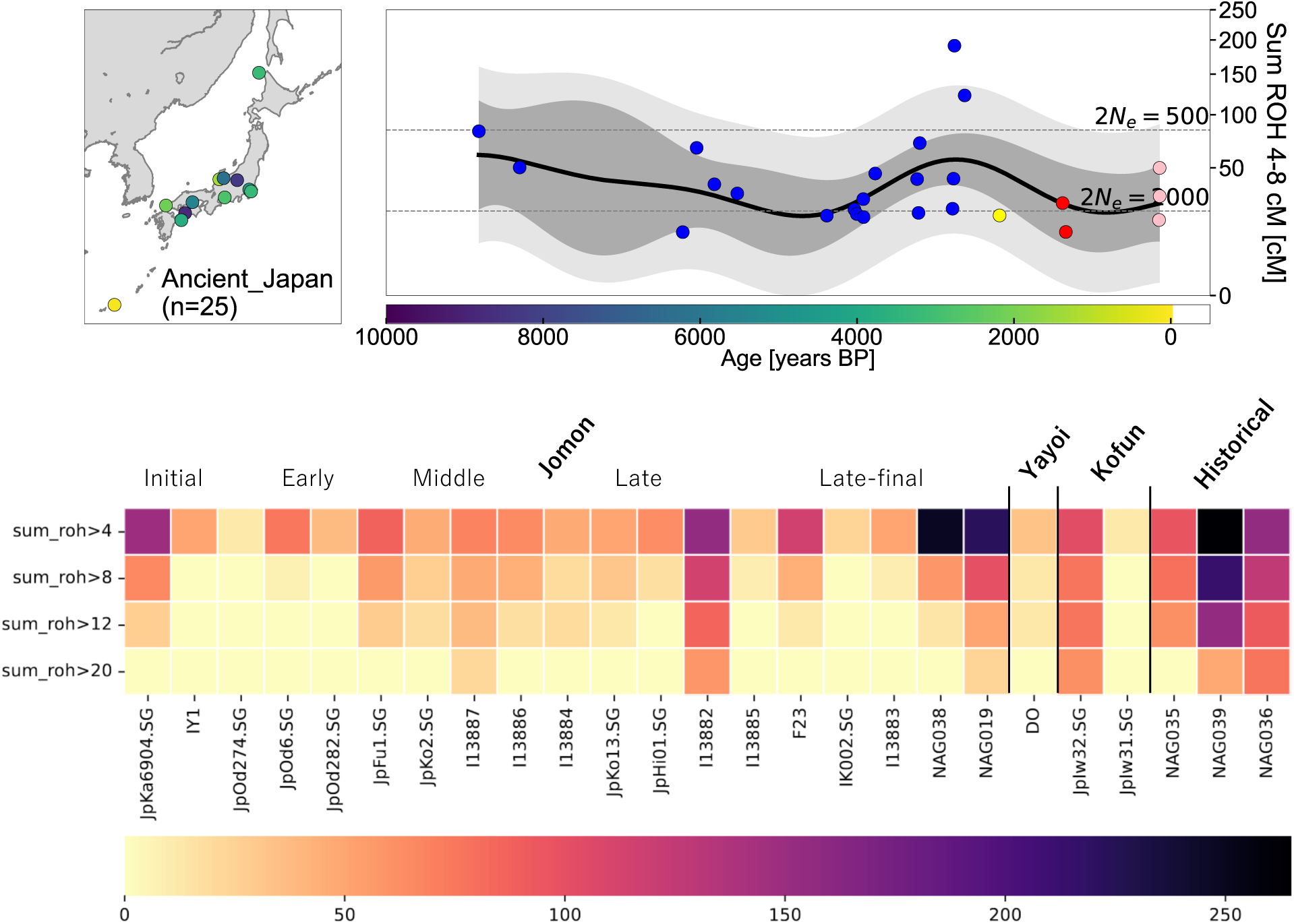
Variations of runs of homozygosity of ancient individuals in the Japanese archipelago. This figure shows ROHs information for each ancient sample from the Japanese archipelago. The upper left figure shows the location of the excavated samples in the Japanese archipelago and their age as indicated by the colors. The plot color on the map indicates the period, with darker blue indicating older and more yellow indicating newer. The upper right figure plots the total number of short ROHs (4-8 cM). Mean estimates were calculated from a Gaussian process model and 95% empirical confidence intervals for both individuals (light grey) and the estimated mean (dark grey). The figure below outlines the total number of ROHs (4-20cM) of various lengths estimated from each individual. The color bars correspond to the size of the number of ROHs. The lighter the beige, the lower the totalized number of ROHs.

### 3.6 Genetic remnants from Jomon and Yayoi lineages

Maximum likelihood trees among populations in East Eurasia suggested that Jomon is an early divergent lineage within East Asians (Figure 9). In addition, migration events from the Jomon to modern Japanese populations have been inferred, supporting the contribution of the Jomon lineage to the Japanese population, as shown in previous studies (Kanzawa-Kiriyama et al., 2019; Gakuhari et al., 2020).

**Figure 9.**
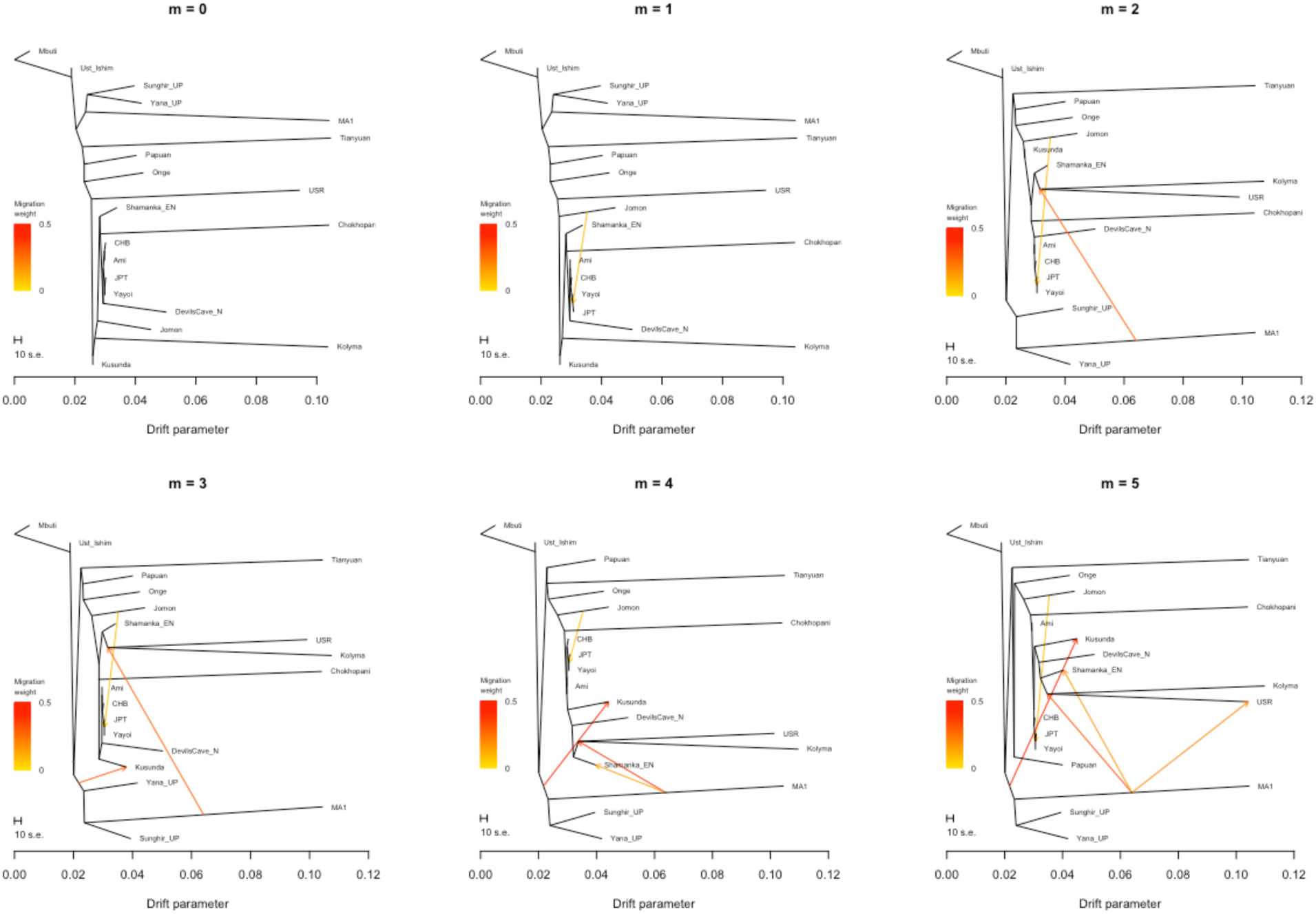
Maximum likelihood phylogenetic tree of populations under migration models (*m*=0 to 5). The tree shows the phylogenetic relationships between modern and ancient Eurasian populations (ancient: Ust_Ishim, Tianyuan, DevilsCave_N, Shamanka_EN, USR, Kolyma, Chokhopani, Yana_UP, MA1, Sunghir_UP, Jomon (IY1), and Yayoi (DO); modern: Papuan, Onge, Ami, CHB, JPT, and Kusunda; outgroup: Mbuti). The tree represents the results from zero to five as the number of migrations (*m*). The migration weight indicates the fraction derived from the source of ancestral migration.

Decays in the weighted linkage disequilibrium (LD) were computed to infer the timing of admixtures targeting present-day Japanese, assuming Jomon (IY1), Yayoi (DO), and Han as the resource populations (Figure 10). The LD decay of IY1 and DO is 72.34 ± 6.82 gen, and that of IY1 and Han is 56.77 ± 3.66 gen, suggesting that the contribution to the modern Japanese population may be more significant from continental immigrants at the Yayoi periods. We calculated the admixture coefficients for present-day Japanese people in different source scenarios. As a result, the nested models with Yayoi-Doigahama (DO), AR_IA, WLR_LN, CHB, YR_MN, Korean, and Upper_YR_IA individuals as one of the resources were not rejected (Table 2, Figure 11).

**Figure 10.**
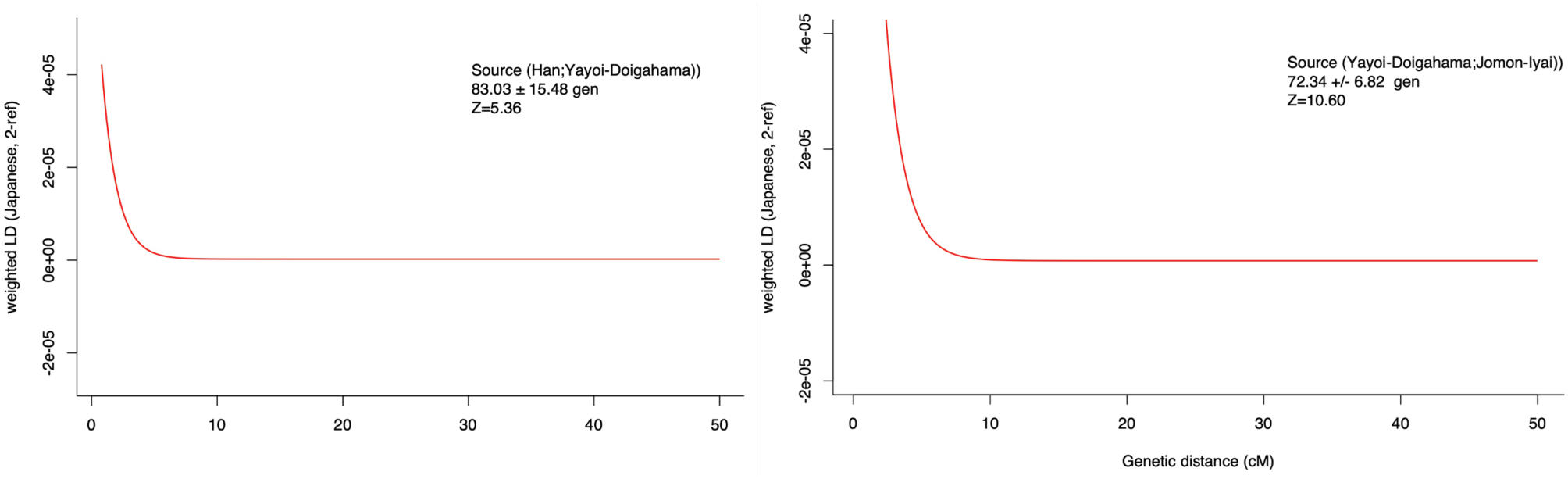
LD decay curves assuming immigrant population in present-day Japanese population.

**Figure 11.**
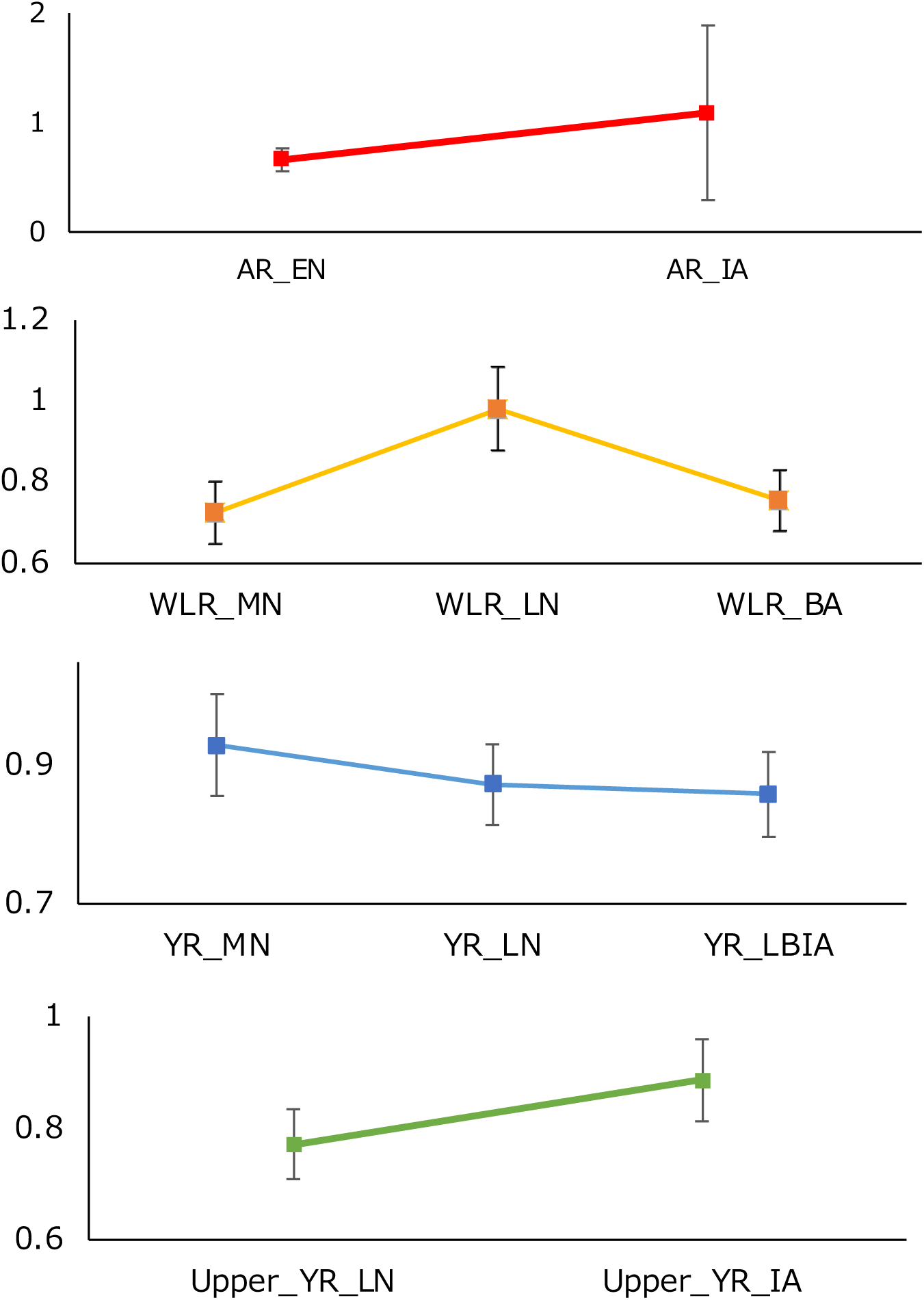
Admixture modeling with ancient Chinese populations. This figure illustrates the coefficient variation by period with ancient Chinese peoples in qpAdm. Yellow River (YR): Middle Neolithic, YR_MN; Late Neolithic, YR_LN; Late Neolithic, Upper_ YR_LN; Late Bronze Age/Iron Age, YR_LBIA; and Iron Age, Upper_YR_IA. West Liao River (WLR): Middle Neolithic, WLR_MN; Late Neolithic, WLR_LN; Bronze Age, WLR_BA. Amur River (AR): Early Neolithic, AR_EN; and Iron Age, AR_IA and AR_Xianbei_IA.

**Table 2.** Inference of admixture proportion in various source scenarios.

| Target | Source X1 | Source X2 | Coefficient |  | std.err | *p*-value for nested model | Rejected |
| --- | --- | --- | --- | --- | --- | --- | --- |
|  |  |  | X1 | X2 |  |  |  |
| Japanese | Yayoi | Jomon | 1.119 | -0.119 | 0.122 | 2.77E-01 | No |
|  | AR_IA | Jomon | 1.09 | -0.09 | 0.795 | 9.17E-01 | No |
|  | WLR_LN | Jomon | 0.981 | 0.019 | 0.103 | 8.82E-01 | No |
|  | CHB | Jomon | 0.970 | 0.030 | 0.057 | 6.40E-01 | No |
|  | YR_MN | Jomon | 0.930 | 0.070 | 0.073 | 3.47E-01 | No |
|  | Korean | Jomon | 0.917 | 0.083 | 0.074 | 2.50E-01 | No |
|  | Upper_YR_IA | Jomon | 0.887 | 0.113 | 0.074 | 1.28E-01 | No |
|  | AR_Xianbei_IA | Jomon | 0.887 | 0.113 | 0.197 | 5.27E-01 | No |
|  | YR_LN | Jomon | 0.873 | 0.127 | 0.059 | 2.92E-02 | Yes |
|  | Kofun | Jomon | 0.864 | 0.136 | 0.064 | 4.52E-02 | Yes |
|  | YR_LBIA | Jomon | 0.859 | 0.141 | 0.061 | 1.90E-02 | Yes |
|  | Upper_YR_LN | Jomon | 0.772 | 0.228 | 0.062 | 2.54E-04 | Yes |
|  | WLR_BA | Jomon | 0.754 | 0.246 | 0.075 | 4.26E-03 | Yes |
|  | Devils'Gate_EN | Jomon | 0.733 | 0.267 | 0.074 | 3.24E-04 | Yes |
|  | WLR_MN | Jomon | 0.724 | 0.276 | 0.077 | 5.50E-04 | Yes |
|  | Boisman_MN | Jomon | 0.716 | 0.284 | 0.063 | 6.26E-07 | Yes |
|  | Miaozigou_MN | Jomon | 0.713 | 0.287 | 0.08 | 2.72E-03 | Yes |
|  | AR_EN | Jomon | 0.665 | 0.335 | 0.103 | 1.14E-03 | Yes |
|  | HMMH_MN | Jomon | 0.647 | 0.353 | 0.062 | 1.01E-06 | Yes |

OG: Mbuti, Ust\_Ishim.DG, Tianyuan, Russia\_Kostenki14, Papuan, ONG, Yana\_UP, Liangdao, Yumin; Jomon: IY1; Yayoi: DO.

### 3.7 Estimation of CNVs in AMY1 gene

Amylases are a group of digestive enzymes found in the pancreatic juice and saliva that convert starch into glucose, maltose, and oligosaccharides by hydrolyzing glycosidic bonds. Variations in the copy number of amylase genes have been observed to correspond to starch consumption in different regional human populations. In this study, the copy number of the AMY1A gene was estimated for two individuals, one from the initial Jomon period (IY1) and the other from the Middle Yayoi period (DO). Both individuals had relatively high copy numbers compared to modern East Asian populations, with IY1 having approximately 9 copies and DO having approximately ten copies of DO (Figure 12). The copy number of AMY1A in La Braña 1, a Mesolithic European hunter-gatherer from the La Braña-Alintero site in Spain, was estimated to be approximately five (Olalde et al., 2014). These results may be related to the different dietary styles of the hunter-gatherer Jomon individuals of IY1 and the Mesolithic Period European hunter-gatherers who lived in the Japanese archipelago.

**Figure 12.**
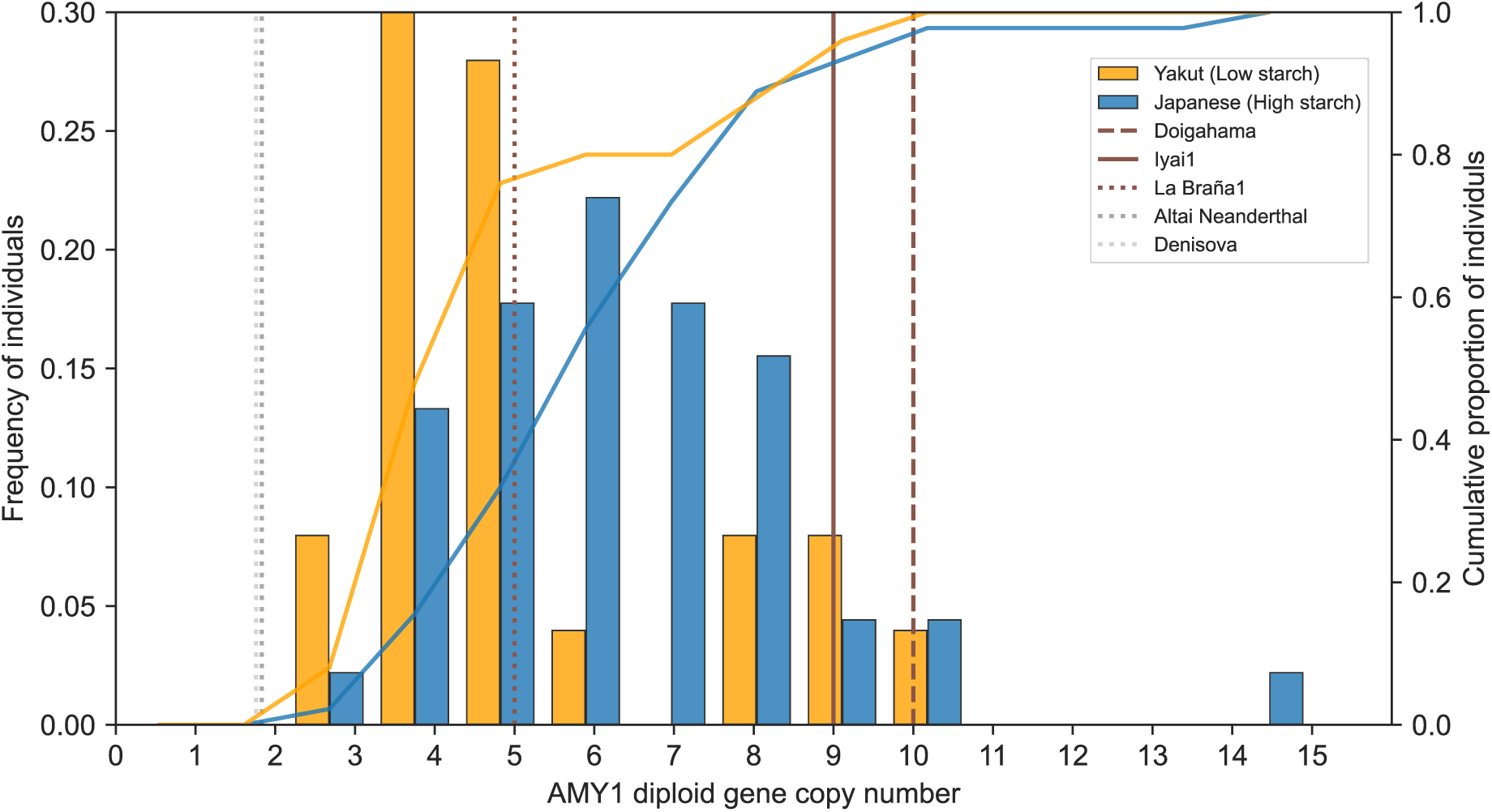
Comparison of copy number of AMY1A gene. This figure outlines the estimated copy number polymorphisms in the AMY1A gene. The distribution of copy number polymorphisms of the gene is compared between low-starch (Yakut) and high-starch (Japanese) dietary populations in Northeast Asia. The lines indicate the cumulative percentage of copy number polymorphisms in each population. The vertical lines show the estimated copy number of AMY1A in each ancient individual.

### 3.8 HLA haplotypes of DO and IY1

Based on 4-digit alleles in HLA, shotgun sequencing data from Yayoi individual DO and Jomon individual IY1 were genotyped using xHLA. Genotyping estimates were obtained for DO and IY1 at six loci (A, B, C, DPB1, DQB1, and DRB1) (Supplementary Table S5). The resulting frequencies of each allele in modern geographic populations were visualized using the Allele Frequency Net Database (Supplementary Figures 14-29). The A*02:06 allele observed in IY1, and the B*15:01 alleles observed in DO were frequently observed in the Japanese Hokkaido Ainu (0.200) and Okinawa Ryukyuans (0.183) (Supplementary Figures S15 and S16). Furthermore, the DPB1*02:01 allele, commonly observed in IY1 and DO, was found at frequencies of 0.282 and 0.279 in modern Japanese and Korean populations, respectively (Supplementary Fig. S22). This allele was observed at a higher frequency (0.359-0.565) in modern Papuans. The DRB1*04:01 allele observed in DO was observed at a higher frequency (0.254-0.361) in Russian Siberian populations (Supplementary Fig. S28).

### 3.8 Archaic ancestry inference in Jomon genome

To estimate the region of introgression from archaic humans in the genome of the initial Jomon individual, we conducted an admixfrog program using the initial Jomon IY1 genome. In this method, the target population was the Jomon, and we assumed states from three resources: the Altai Neanderthal and Denisova genomes were used as archaic resources, and the 1 KGP/HGDP Sub-Saharan Africans were used as present-day human resources. Here, we extracted segments with a posterior probability of Neanderthals or Denisova greater than 95%. Finally, putative archaic (posterior probe. of Denisova: 0.970874) segment was identified on chromosome IY1 (Supplementary Figure S6; Table S1). This segment is located in the OCA2 gene, which has been previously reported to have the strongest influence on skin type in modern Japanese people (Shido et al., 2019).

## 4. Discussion

This study presents low-contamination and high-coverage ancient human genomes from the Japanese Archipelago. These high-quality ancient human genomes represent milestones in our understanding of the Jomon and Yayoi people. Ancient genome research has increasingly included population-scale analyses, using multiple samples from various regions. These studies provide valuable clues for understanding the relationships between populations and their origins across a broad area. However, many samples have less than 1-fold genome coverage, which limits the reliability and applicability of analytical methods. Previous studies have reported >45-fold high-coverage ancient genomes, mainly from high-latitude regions (Prüfer et al., 2014; Günther et al., 2018; Kanzawa-Kiriyama et al., 2019), which serve as benchmarks for discussing population and phylogenetic history. However, the warm and humid climate of Asia, including mainland Japan, makes it particularly challenging to obtain high-quality ancient genomes. Here, we obtained high-coverage whole genomes of the Initial Jomon and migratory Yayoi people from mainland Japan. One reason for the high coverage of these two samples could be the geological and environmental conditions under which the samples were buried. Unearthed human remains from other initial Jomon sites are rare, even in the Japanese archipelago, and many are not well preserved. However, the burial remains unearthed at the Iyai site, although more than 8,000 years old, are mostly well-preserved, allowing DNA analysis with high reliability. The main reason for the good preservation of human remains is the properties of the ash-rich layers of soil that were piled up inside the rock shelter, which were free from rainwater. The layers in which the human remains were buried contained a high proportion of calcite derived from wood ash, which became slightly alkaline owing to the action of CaCO_3_. Consequently, these chemical components of the soil protected the bone tissue and maintained the human remains in good condition. In addition, the preservation of the human remains found at the Doigahama site was extremely good, with complete human remains recovered from the undisturbed burial pits. Even when human remains are disturbed, preserving the remaining bones is extremely good. The cemetery at the Doigahama site is located on a dune consisting of a layer of wind-blown sand. This sand layer is composed of numerous shell fragments and mineral particles. The calcium ions released from the shells by rainwater have an acid-reducing effect. In addition, the human remains are located approximately 1.5 to 2.0 meters below the surface of the site, where moderate humidity is always maintained. The large number of shell particles (calcium) and moderate humidity preserve human bones and organic materials.

The high-coverage genomes of mainland and initial-Jomon individuals, which may have diverged early in the East Asian lineage, and the Yayoi individual, which migration events from Eurasia may have influenced, offer new insights into the genetic context of transition in this region. These two high-coverage genomes from mainland Japan facilitated the acquisition of individual diploid sequences and permitted the estimation of past population dynamics. Our population dynamics estimates highlight ancestral contrasts in demographic history during the Initial-Jomon and Middle Yayoi periods (Figures 7A and 7B). The differences between the two populations became clearer, particularly in the changes after the maximum cold period (Figure 7B). Estimates from the Initial-Jomon individuals suggest that they experienced a drastic population decline after the LGM. In contrast, Yayoi population estimates show a gradual population growth trend after 10,000 years. This may reflect environmental changes and differences in the lifestyles of the ancestral populations. The Jomon people, who have persisted in the archipelago for a long time, are thought to have had diverse lifestyles and diets in each region, and their hunter-gatherer-fisher lifestyles could have been affected by changes in the natural environment. In addition, in the cross-period comparison of ROH against ancient genome data excavated from the Japanese archipelago, several Jomon individuals showed high ROHs from the Late to Late-final Jomon period. One potential factor contributing to the observed increase in ROHs may be the substantial reduction in population size. Two possible reasons can explain this event, interaction with other regions and significant environmental shifts. This finding was consistent with the decrease in population during the Late Jomon period, which was inferred from the Y-chromosome of the present-day Japanese (Watanabe et al., 2019). A previous archaeological study on the number of Jomon sites indicated a relationship between the decrease in temperature and population size during the Late to Late-final Jomon period (Koyama 1979). A study of paleotemperatures also showed a decrease in temperature during this period, as reported by Kawahata et al. (2017), who reconstructed paleotemperatures using alkenone sea surface temperature (SST) measurements from coastal sedimentary cores. However, the demographics of the same period require sufficient verification based on regional differences, owing to the long period duration, customs, and interactions within each archaeological site. In the future, conducting comprehensive genomic data analysis of Jomon period individuals from various regions within the same timeframe will be essential.

AMY1 copy number is well known to correlate with amylase protein levels and enzyme activity, although some argue that CNV plays a limited role in its functions (Carpenter et al. 2017). AMY gene was strongly selected in humans as early as the Middle Pleistocene (Inchley et al. 2016). In contrast, no estimated increase in AMY1 copy number has been observed in archaic humans, such as Neanderthals and Denisovans (Perry et al., 2015). Directive evidence of CNVs in ancient humans suggests that our ancestors evolved specific adaptations to digest starch-rich foods. However, estimating copy number variations (CNVs) in ancient genome sequencing is challenging because of insufficient coverage across the human genome. Our high-coverage data enabled us to detect copy number variations in the human genome. This study focused on copy number polymorphisms in AMY1, a salivary amylase gene that varies across regional human populations (Groot et al., 1989; Iafrate et al., 2004). Previous research on AMY1 has indicated that copy number polymorphisms are associated with human dietary habits, particularly among populations consuming a high-starch diet, which tend to have higher copy numbers (Perry et al., 2007). Genome analysis of ancient European hunter-gatherers suggested that the low copy number of AMY1 may be related to their poor dietary style of digesting starch (Olalde et al., 2014). Our findings imply that populations of IY1 and DO have higher copy numbers than hunter-gatherers in Europe. The Jomon people are thought to have followed a hunter-fisher-gatherer way of life for a long time (Koyama et al., 1979) and may have consumed starch-rich nuts and grains. The potential cause of the copy number polymorphism of AMY1 in the initial Jomon individual may have been their dietary habits. However, the copy number of Yayoi DO was estimated to be even higher than that of Jomon DOs. In light of the differing distributions of AMY1 copy number variations in modern human populations consuming high– and low-starch diets, our inferences regarding ancient genomic data do not definitively preclude the possibility that individuals residing in the Japanese Archipelago during a specific period rely on a starch-rich diet. The Yayoi period has been proposed to have been influenced by migrants thought to have introduced a paddy field rice farming culture to the island. The direct observation of CNVs in ancient samples, as in our approach, is useful for studying variations in AMY gene copy numbers distribution among regional populations.

The impact of continental arrival events on the people of the Japanese Archipelago has long been widely recognized as significant. The lack of high-quality ancient genomes and migratory lineages of individuals in the surrounding areas has hindered the adequate consideration of migrant population resources. We used the Japanese population as our target population to examine the timing of hybridization with resource populations. Using the Jomon, Yayoi, and Han Chinese as our resource populations, our analysis estimated the timing of the Yayoi and Jomon admixture between 65.52 and 79.16 generations (Figure 10). Assuming a generation time of 25 years, we estimate this result to be between 1,638 and 1,979 years ago, which corresponds to the Yayoi period, the period in which migration is assumed to have occurred. In addition, the timing of admixture between the present Han Chinese and Yayoi populations was between 67.55 and 98.51 generations (Figure 10), suggesting that admixture on the continental side occurred earlier than that in the Jomon population. Furthermore, we performed admixture, *f3* and *f4* statistics on high-coverage Jomon (IY1) and Yayoi (DO) individuals to explore the origin of the non-Jomon genetic component of the Yayoi people (Figure 4, 5, and 6). Admixture analysis focused on components observed in the Yayoi individual but not in Jomon people and examined their geographical distribution (Figure 4). Interestingly, this component has often been observed in ancient Central Eurasian individuals from the Ust’-Belaya burial ground near Lake Baikal (Flegontov et al., 2019) and Mongolia (Wang et al., 2021) (Figures 3 and 4). However, this proportion decreased in Mongolia after 500 BCE (Figure 3, Supplementary Figure S10).

Yayoi individual (DO) had a significant genetic affinity for Middle Neolithic individuals from Inner Mongolia, Miaozigou (Miaozigou_MN), and the Haminmangha site (HMMH_MN). Furthermore, DO also showed significant affinities with early Neolithic individuals (Xiaojingshan and Boshan) in Shandong, China. In present-day Northeast Asia, Daur people also showed a significant *f4* value with DO (Figure 5). These results suggest that Eurasian coastal regions and Central Asia or Northeast Asia could influence Yayoi-related ancestral populations.

The results of our analysis also highlight differences in the influence of the Far East region of Eurasia on the Yayoi and Kofun individuals. The *f3*-statistics of the Yayoi individual provide stronger support for an affinity with Devil’sGate and Boisman individuals in the Far East region than with those of the Kofun period (Figure 6). These results suggest that ancestral genetic contexts differ between Yayoi and Kofun and imply that the Yayoi individual are more strongly influenced by the Far East region. Because the spread of paddy rice cultivation to the Japanese archipelago may have been linked to continental migration, the availability of ancient genome data from the lower Yangtze River region in the future may provide further clarity regarding the dissemination of paddy rice cultivation.

Wang et al. (2021) suggested that hunter-gatherers from the Baikal region and the Amur River basin, including Boisman, possess a presumed Mongolian Neolithic-related ancestry. Our investigation does not preclude the possibility that the Yayoi individual, who exhibited a high degree of genetic similarity to these two populations, were also influenced by Mongolian Neolithic-related ancestors. In other words, one of the ancestral components of the Yayoi people may have been derived from the population that brought paddy rice cultivation to Japan from East Eurasia and Central Eurasian Neolithic-related ancestry.

In our study of the high-coverage Jomon genome, we identified components that were not present in the previously published Jomon genomes (Figure 3). The ancestral component may have been overlooked or underestimated in the low-coverage ancient genome data. Our analysis reassesses the dual structure model of the Japanese formation proposed by Hanihara (1991). A recent study proposed a tripartite origin for Japan, including the impact of the Kofun Period (Cook et al., 2021). They concluded that it was Kofun Period when the ancestral components of present-day Japanese were established. However, the genomic coverage of the Yayoi individual used in this previous study was limited (Supplementary Table S3), and their origin should be discussed carefully. In our analysis, the individual from the mid-Yayoi period showed components observed in present-day Japanese, confirming the continuity of common ancestral components from the Yayoi period (which is earlier than Kofun period) to present-day Japanese, as shown by Kim et al., (under revision). Our results emphasize that the growth of high-coverage ancient genomic data could shed light on the origins of the human population, which would not limited to the Japanese Archipelago.

In general, our results emphasize the existence of a population with a continental component in the middle Yayoi period that differed significantly in genetic composition from the previously indigenous Jomon period individuals. In contrast, our admixture analysis also found a common genetic component between Jomon and Yayoi individuals. This common component was observed in the Papua New Guinean and Vanuatu people and may have been derived from an old Asian lineage. The results of our admixture modeling also suggest that the common ancestral resources of the Yayoi and Jomon people may be related to early divergent old Asian lineages (Supplementary Figure S12, S13).

During the out-of-Africa process in Homo sapiens, they encountered and interbred with archaic hominins, such as Neanderthals and Denisovans (Green et al., 2010; Reich et al., 2010). Archaic introgressed DNA segments have also been suggested to contribute to human phenotypic traits (Huerta-Sanchez et al., 2014; Racimo et al., 2017; Dannemann et al., 2017). However, our knowledge of archaic DNA segments in ancient humans in East Asia, including Japan, is still limited. Here, we found a putative archaic introgressed segment in OCA2 on human chromosome 15q11.2-q12 in the initial Jomon individual (Supplementary Figure S30; Supplementary Table S4). This gene plays an important role in skin, hair, and eye color formation and is known to strongly influence skin type, especially in Japanese people (Shido et al., 2019). However, no similar segments were found in the Yayoi or modern Japanese individuals. Interestingly, a previous study reported that Neanderthal-derived introgression regions were more frequent in OCA2 in East Asian populations than in other regional populations (Gittelman et al., 2016). Possibly, the OCA2 region, where adaptive introgression occurred in ancient East Asia, may have had resources of other archaic origins as well as Neanderthals. Our high-coverage ancient genome resource offers a promising opportunity for gaining new insights into the pursuit of remnants and interactions among prehistoric populations in the East Eurasian region.

## Acknowledgements

This study was approved by the Ethics Committee of Toho University School of Medicine (A23103_A20110_A18099_A18056). We would like to express our gratitude for the contributions of Michiko Hayashi to the experiments and the efforts of Yusuke Takemura to investigate the published genome data. We would like to thank Editage (www.editage.jp) for English language editing.

