## Supplementary Figures for "High-coverage genome sequencing of Yayoi and Jomon individuals shed light on prehistoric human population history in East Eurasian"

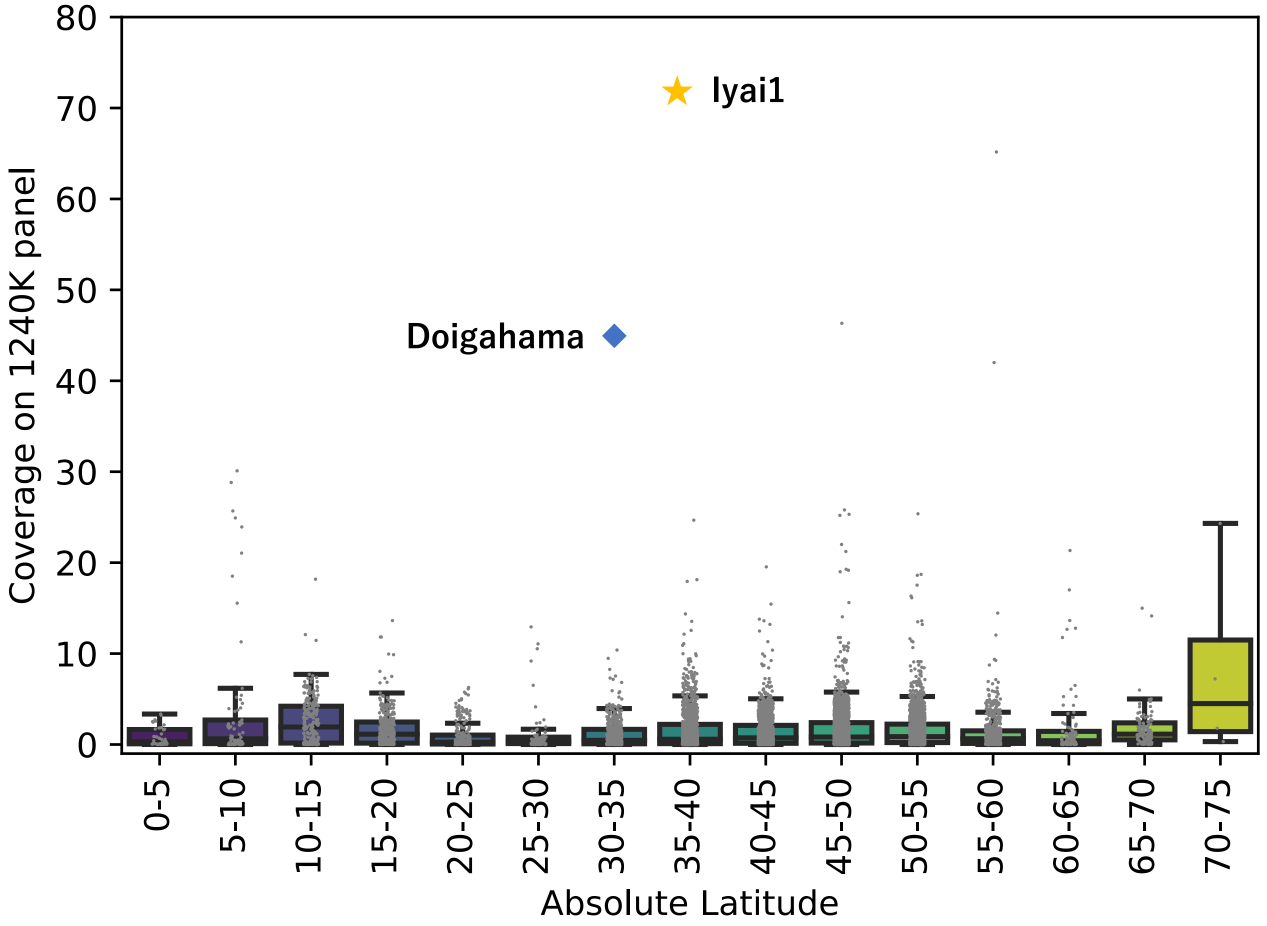


**Figure S1**. Comparison of SNP coverages across published ancient samples.

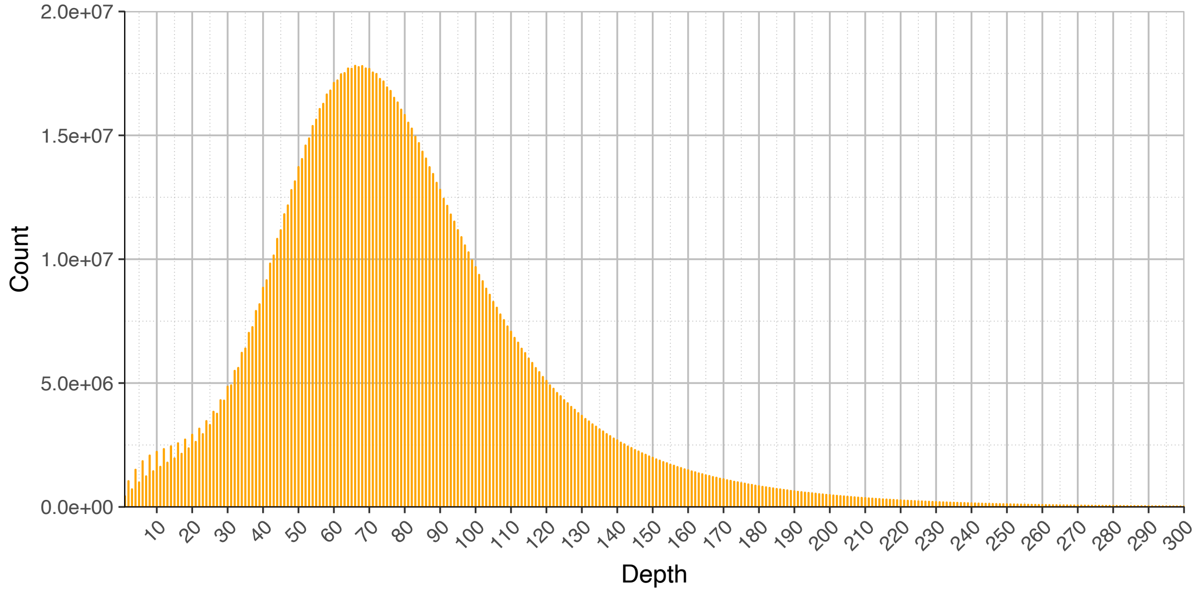


**Figure S2.** Distribution of sequencing read coverage of the Initial-Jomon sample IY1.

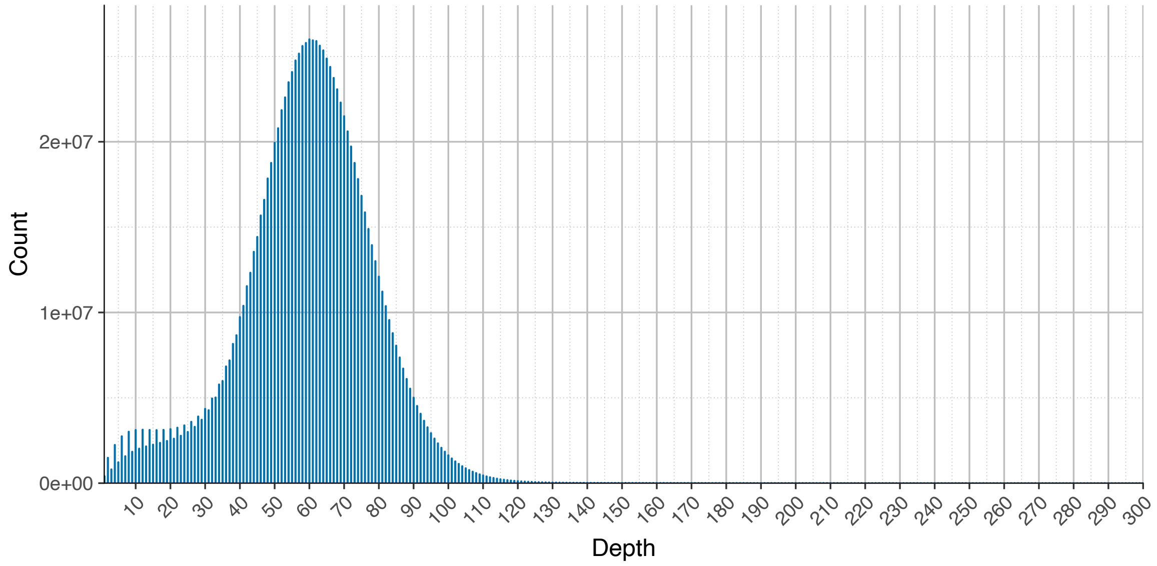


**Figure S3**. Distribution of sequencing read coverage of the Middle-Yayoi sample DO.

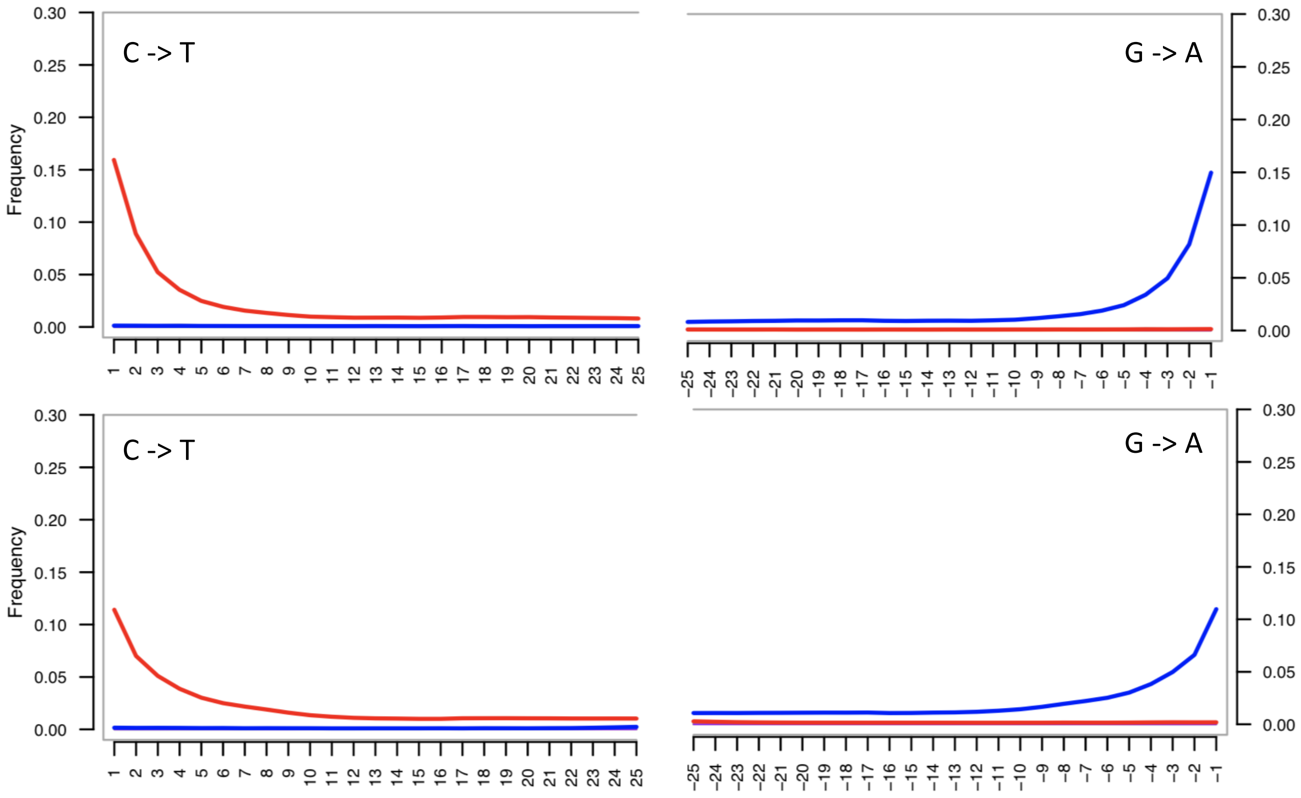


**Figure S4**. Deamination patterns on mapped reads from IY1 (upper) and DO (lower).

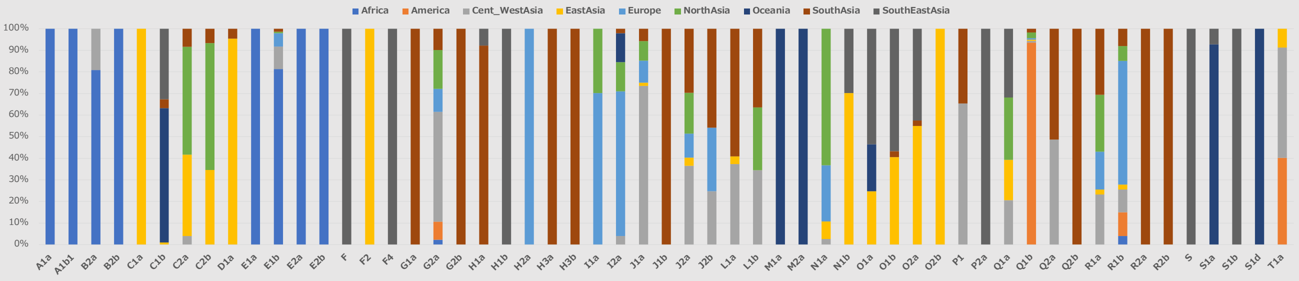


**Figure S5**. Proportion of Y-chromosome haplogroups in the worldwide populations (n=1,195).

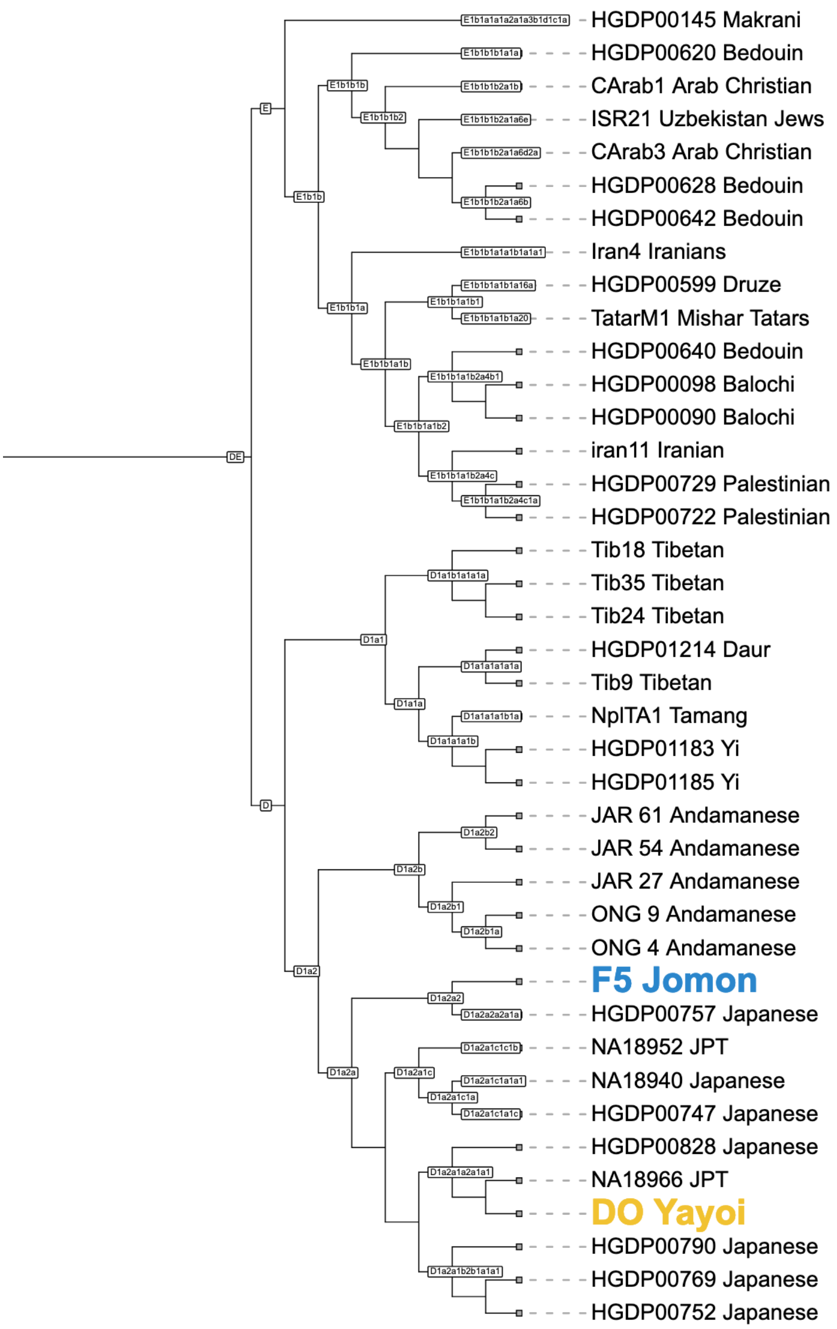


**Figure S6**. Phylogenetic relationships based on Y-chromosome genome wide SNPs.

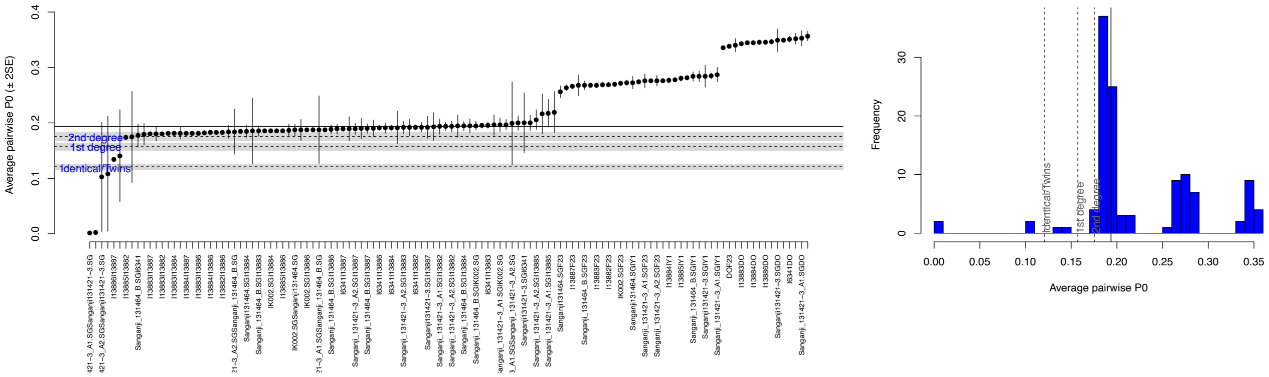


**Figure S7**. A kinship analysis in published ancient Japanese samples merged with newly sequenced samples (DO and IY1).

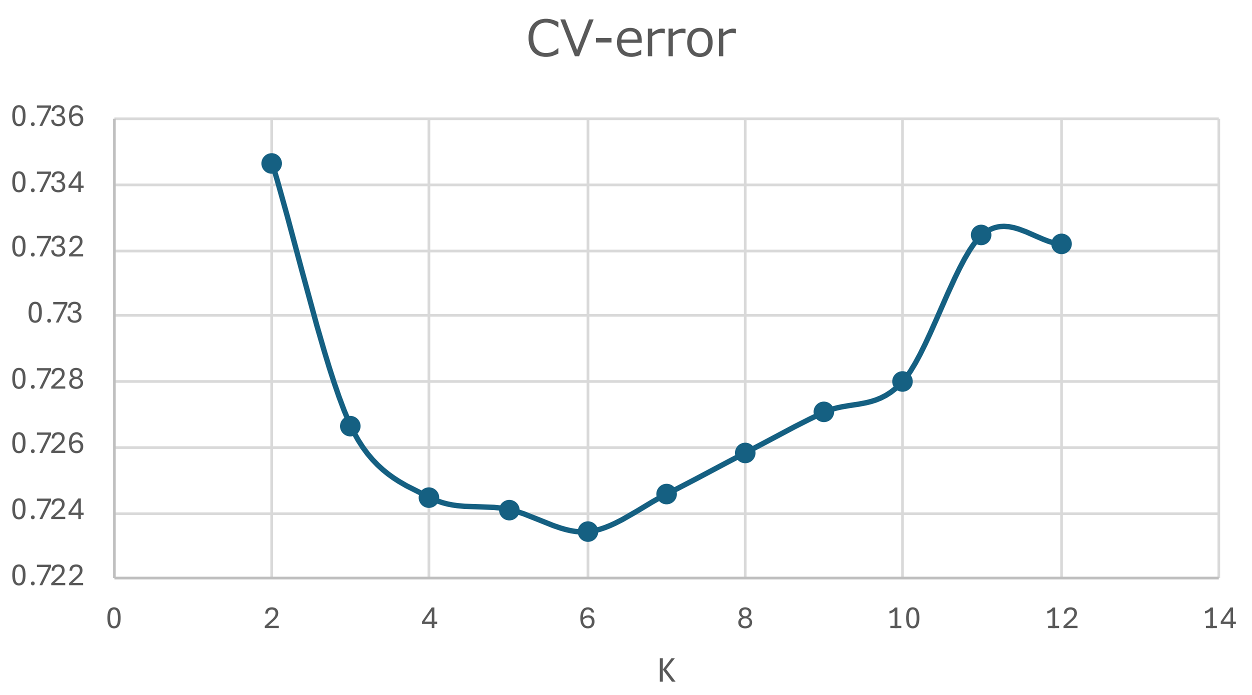


**Figure S8**. Cross-validation errors in Admixture analysis (from *K*=2 to 12). The lowest cross-validation error was shown when the ancestral component was *K*=6.

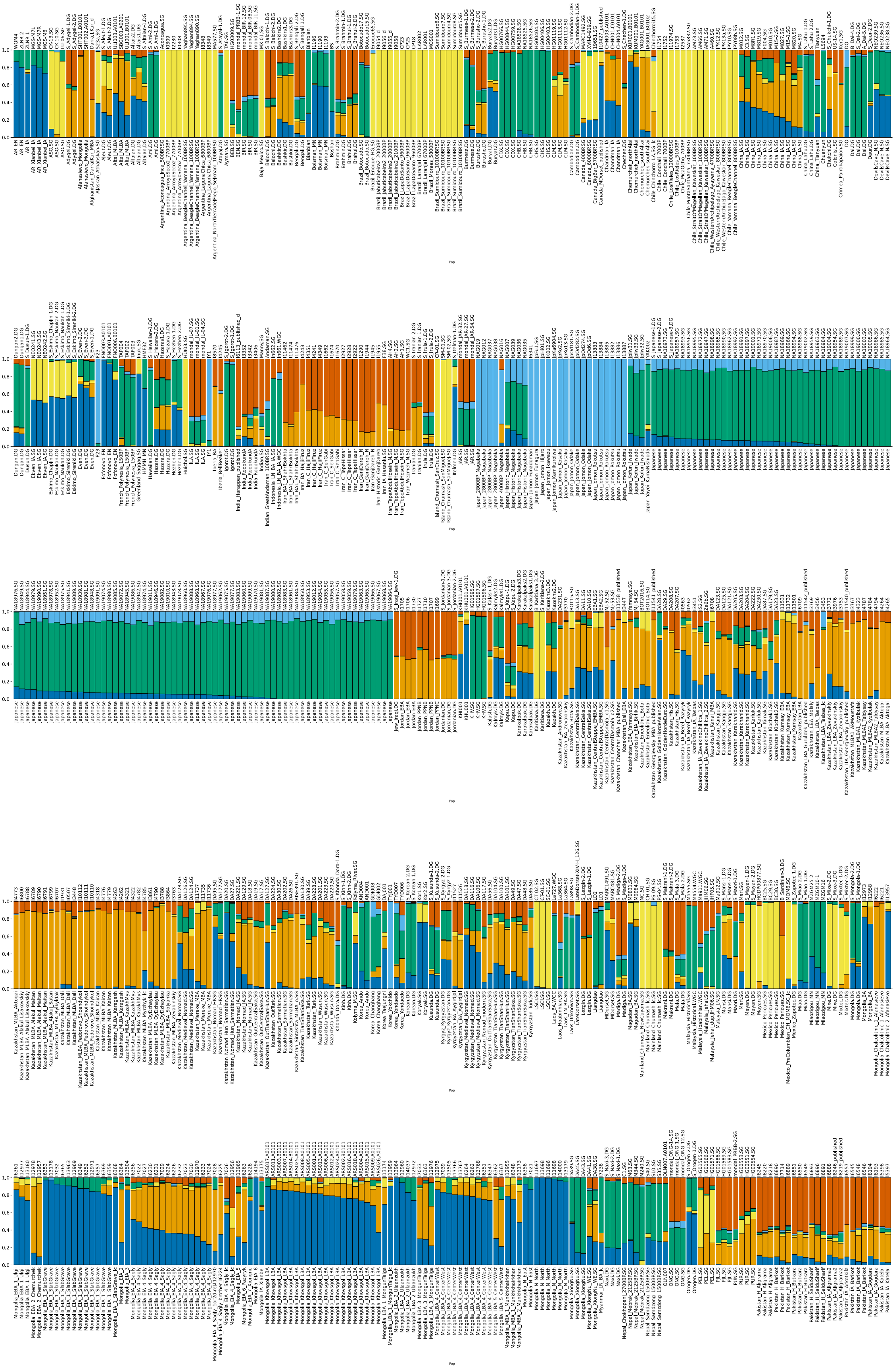

**Figure S9** (page1/2). The results of the Admixture analysis under *K*=6 with the lowest cross-validation error. Each bar plot shows the percentage of ancestral components.

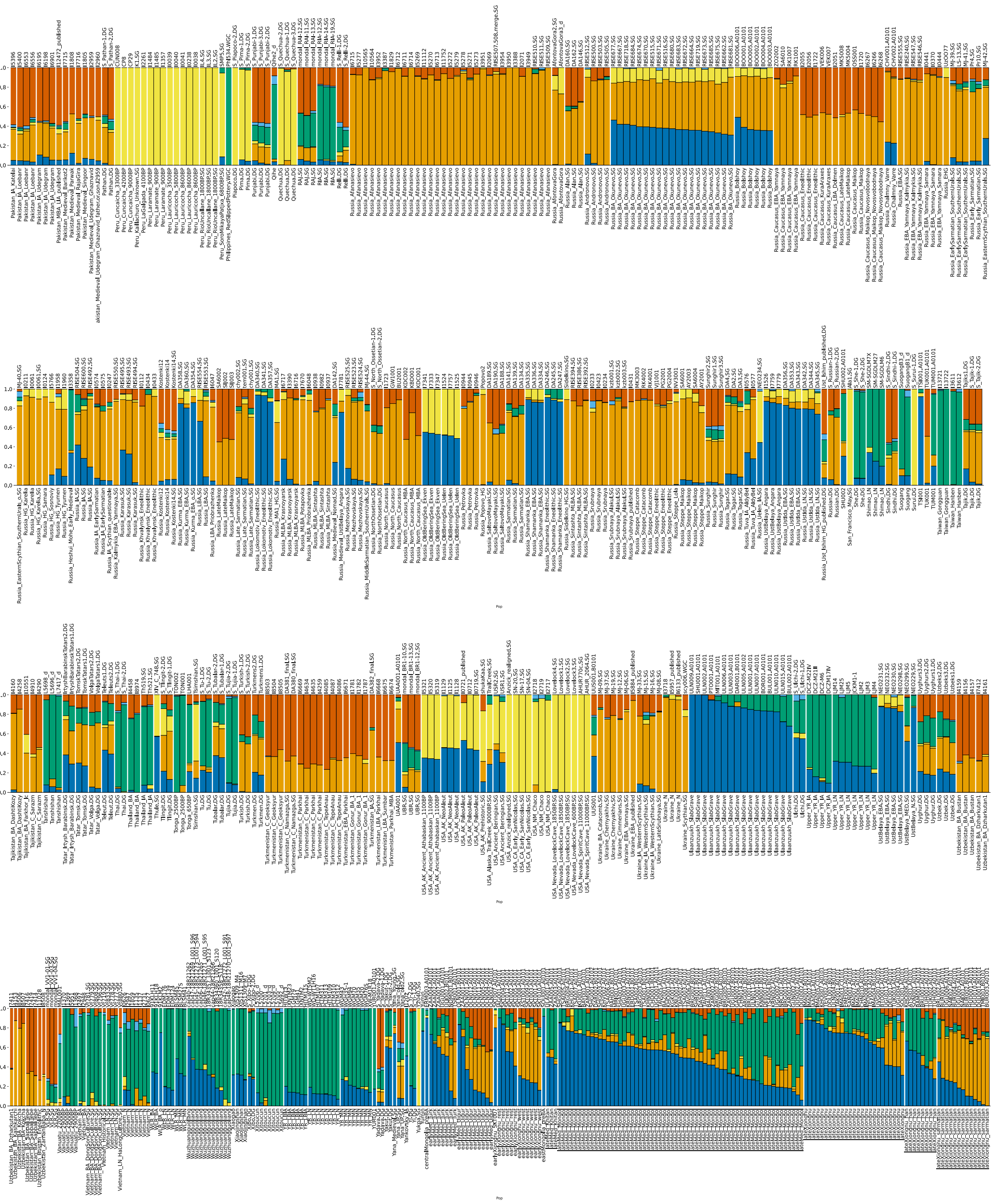

**Figure S9** (page2/2). The results of the Admixture analysis under *K*=6 with the lowest cross-validation error. Each bar plot shows the percentage of ancestral components.

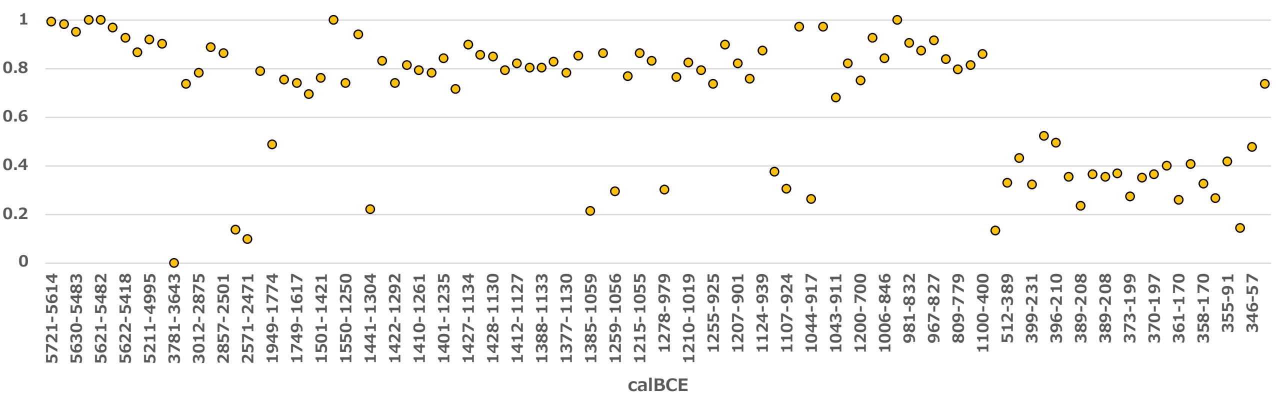


**Figure S10**. The frequency of the ancestral component (P1) observed among ancient Mongolia samples. The horizontal axis shows the calibrated radiocarbon date (calBCE) of the sample and the frequency of the ancestral component (P1) of each individual on the horizontal axis.

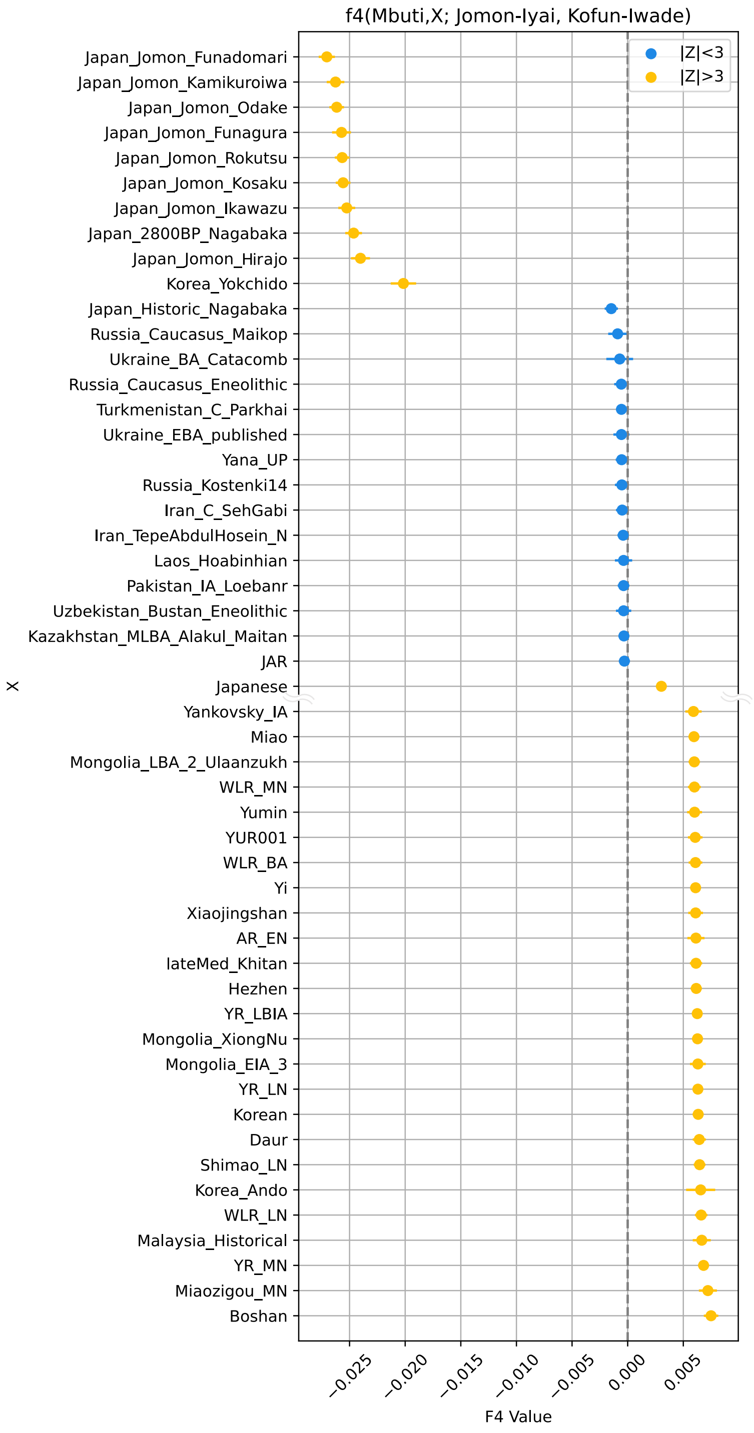


**Figure S11.** Results of *f4*-statistics highlighting the shared genetic drift between Initial-Jomon (Jomon-Iyai) and Kofun (Kofun-Iwade). In *f4*(Mbuti; X, Jomon-Iyai, Kofun), the top 50 individuals/populations were selected from those with the highest absolute *f4* values. Error bars indicate the standard deviation; absolute values of *Z* greater than 3 are shown in yellow and values less than 3 in blue. Negative values indicate more shared genetic drift with Jomon-Iyai and positive values with Kofun.

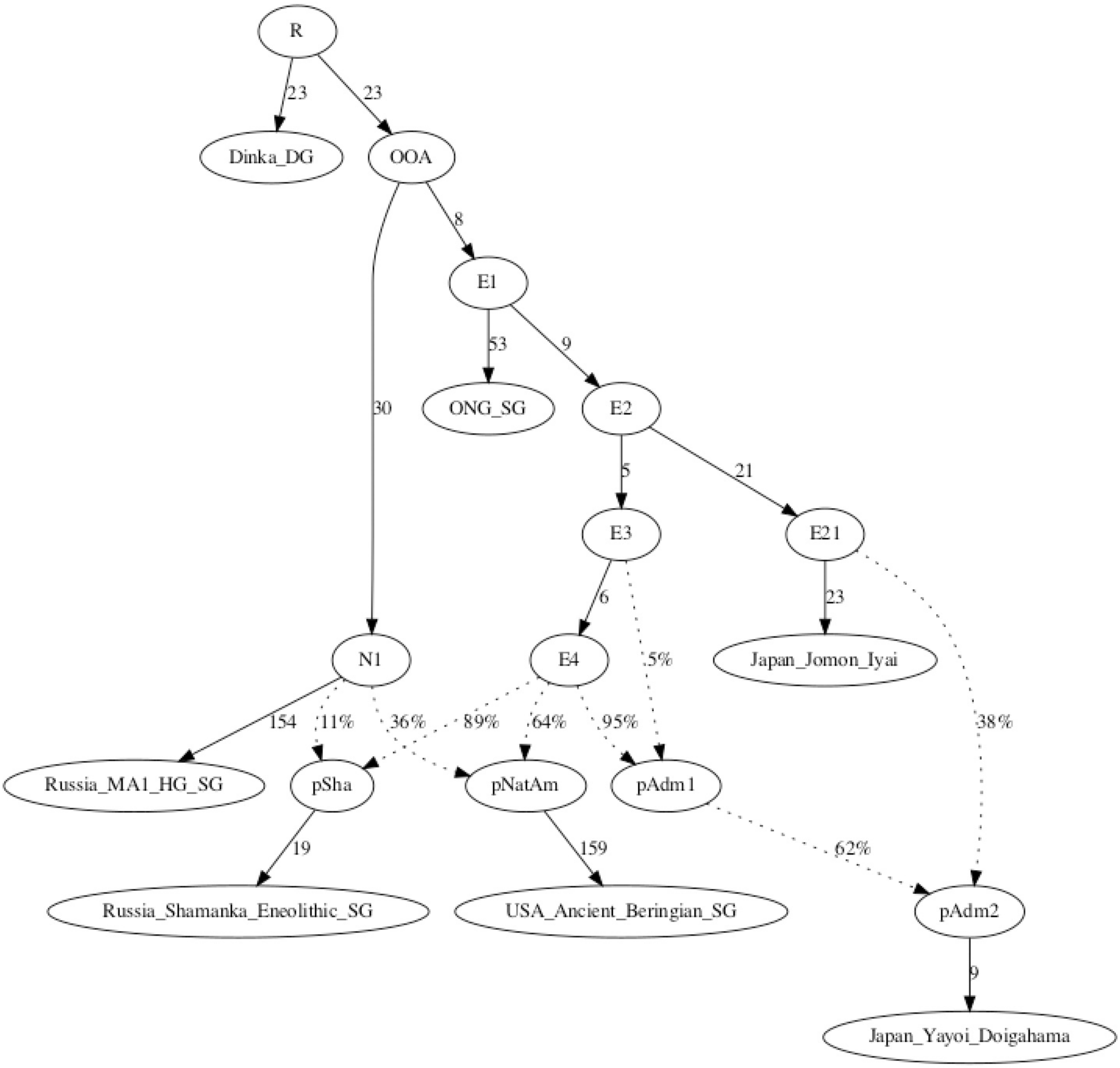


**Figure S12**. An admixture modeling under possible ancestral sources (Z=-2.39).

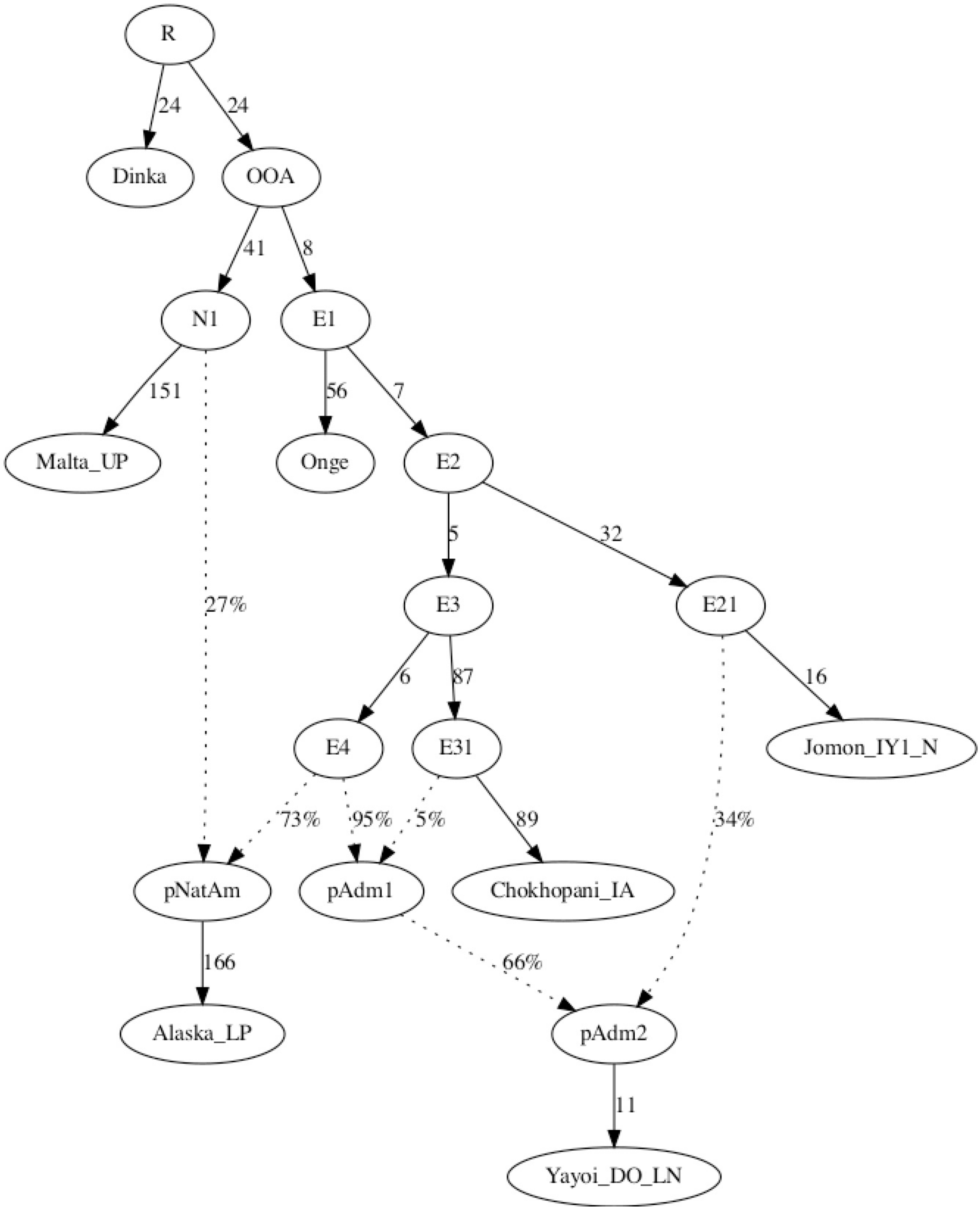


**Figure S13**. An admixture modeling under possible ancestral sources (Z=-3.120).

| Population | Phenotype Frequency (%) | Allele Frequency (in_decimals) | Sample Size |
|---|---|---|---|
| Mexico Sonora Seri | | 0.5450 | 34 |
| Peru Titikaka Lake Uro | | 0.5000 | 105 |
| Peru Titikaka Lake Uros | | 0.5000 | 105 |
| USA Arizona Gila River Amerindian | | 0.4740 | 492 |
| USA Arizona Pima | | 0.4360 | 100 |
| Malaysia Kedah Kensiu | 57.0 | 0.4050 | 21 |
| Argentina Gran Chaco Western Toba Pilaga | 60.0 | 0.4000 | 19 |
| Finland | | 0.3440 | 91 |
| Mexico Mestizo | 56.1 | 0.3410 | 41 |
| KOS Kosovo | 52.4 | 0.3186 | 124 |

**Figure S14**. Screenshot of A*02:01 allele frequencies in modern human populations (high frequency top10 population data).

| Population | Phenotype Frequency (%) | Allele Frequency (in_decimals) | Sample Size |
|---|---|---|---|
| Mexico Oaxaca Mixe | | 0.3490 | 55 |
| New Zealand Maori with Full Ancestry | 52.2 | 0.2940 | 46 |
| Mexico Oaxaca Zapotec | | 0.2610 | 90 |
| New Zealand Maori with Admixed History | 41.0 | 0.2240 | 105 |
| Pakistan Kalash | | 0.2160 | 69 |
| Mexico Oaxaca Mixtec | | 0.2160 | 103 |
| Japan Hokkaido Ainu | | 0.2000 | 50 |
| Japan Okinawa Ryukyuan | | 0.1830 | 143 |
| Mexico Oaxaca Jamiltepec Mixtec | | 0.1670 | 96 |
| Mexico Mixtec | | 0.1650 | 97 |

**Figure S15.** Screenshot of A*02:06 allele frequencies in modern human populations (high frequency top10 population data).

| Population | Phenotype Frequency (%) | Allele Frequency (in_decimals) | Sample Size |
|---|---|---|---|
| China Beijing pop 2 | | 0.4007 | 826 |
| Japan Hokkaido Ainu | | 0.2900 | 50 |
| Sweden Northern Sami | | 0.1500 | 154 |
| Mexico Chihuahua Tarahumara | | 0.1480 | 44 |
| Sweden Southern Sami | | 0.1400 | 130 |
| Mexico Tamaulipas, Ciudad Victoria | | 0.1304 | 23 |
| Colombia North Wiwa El Encanto | | 0.1250 | 52 |
| China Tibet Region Tibetan | | 0.1230 | 158 |
| Netherlands UMCU | 23.4 | 0.1172 | 64 |
| China Shanxi HIV negative | | 0.1140 | 22 |

**Figure S16.** Screenshot of B*15:01 allele frequencies in modern human populations (high frequency top10 population data).

| Population | Phenotype Frequency (%) | Allele Frequency (in_decimals) | Sample Size |
|---|---|---|---|
| Bulgaria Romani | | 0.2730 | 13 |
| India West Bhil | | 0.1900 | 50 |
| India Mumbai Maratha | | 0.1790 | 91 |
| India Khandesh Region Pawra | | 0.1700 | 50 |
| India North pop 2 | | 0.1540 | 72 |
| South Africa Natal Tamil | | 0.1430 | 51 |
| India Andhra Pradesh Golla | | 0.1300 | 111 |
| Singapore SGVP. Indian INS | | 0.1250 | 86 |
| India Tamil Nadu | | 0.1116 | 2492 |
| Iran Baloch | | 0.1110 | 100 |

**Figure S17.** Screenshot of B*40:06 allele frequencies in modern human populations (high frequency top10 population data).

| Population | Phenotype Frequency (%) | Allele Frequency (in_decimals) | Sample Size |
|---|---|---|---|
| Sweden Northern Sami | | 0.2000 | 154 |
| USA Alaska Yupik | | 0.1150 | 252 |
| Sweden Southern Sami | | 0.1050 | 130 |
| USA NMDP Alaska Native or Aleut | | 0.0890 | 1376 |
| USA North American Native | | 0.0860 | 187 |
| USA Arizona Pima | | 0.0790 | 100 |
| USA South Dakota Lakota Sioux | | 0.0740 | 302 |
| Belgium | 14.3 | 0.0710 | 99 |
| USA Arizona Gila River Amerindian | | 0.0620 | 492 |
| Finland | | 0.0610 | 91 |

**Figure S18.** Screenshot of B*27:05 allele frequencies in modern human populations (high frequency top10 population data).

| Population | Phenotype Frequency (%) | Allele Frequency (in_decimals) | Sample Size |
|---|---|---|---|
| Colombia North Chimila Amerindians | | 0.4680 | 47 |
| Australia Kimberly Aborigine | | 0.2860 | 41 |
| Bulgaria Romani | | 0.2730 | 13 |
| Papua New Guinea Wosera Abelam | | 0.2320 | 131 |
| Australia Yuendumu Aborigine | | 0.2030 | 191 |
| Georgia Tibilisi Kurd | | 0.1550 | 31 |
| Spain Andalusia Romani | | 0.1470 | 99 |
| Australia Groote Eylandt Aborigine | | 0.1370 | 75 |
| New Zealand Polynesians with Admixed History | 18.5 | 0.1110 | 27 |
| USA NMDP South Asian Indian | | 0.1077 | 185391 |

**Figure S19.** Screenshot of C*15:02 allele frequencies in modern human populations (high frequency top10 population data).

| Population | Phenotype Frequency (%) | Allele Frequency (in_decimals) | Sample Size |
|---|---|---|---|
| Mexico Chihuahua Tarahumara | | 0.3980 | 44 |
| Brazil Terena | 54.0 | 0.3510 | 60 |
| Russia Sakhalin Island Nivkhi | | 0.3210 | 53 |
| Mexico Chiapas Lacandon Mayans | | 0.2798 | 218 |
| China Guizhou Province Miao pop 2 | | 0.2350 | 85 |
| China Guizhou Province Bouyei | | 0.2230 | 109 |
| Papua New Guinea Wosera Abelam | | 0.2230 | 131 |
| Mexico Oaxaca Mixe | | 0.2170 | 55 |
| USA NMDP Alaska Native or Aleut | | 0.2060 | 1376 |
| USA Arizona Gila River Amerindian | | 0.1950 | 492 |

**Figure S20.** Screenshot of C*03:04 allele frequencies in modern human populations (high frequency top10 population data).

| Population | Phenotype Frequency (%) | Allele Frequency (in_decimals) | Sample Size |
|---|---|---|---|
| Cameroon Baka Pygmy | | 0.2500 | 10 |
| Russia North Ossetian | 40.2 | 0.2402 | 127 |
| USA San Francisco Caucasian | | 0.2130 | 220 |
| Ireland South | 39.2 | 0.2120 | 250 |
| Germany Essen | 36.2 | 0.2090 | 174 |
| Italy South | | 0.2060 | 141 |
| Cameroon Sawa | | 0.1920 | 13 |
| England North West | 35.2 | 0.1900 | 298 |
| Thailand Northeast pop 2 | | 0.1840 | 400 |
| Germany DKMS - Italy minority | | 0.1826 | 1159 |

**Figure S21.** Screenshot of C*07:01 allele frequencies in modern human populations (high frequency top10 population data).

| Population | Phenotype Frequency (%) | Allele Frequency (in_decimals) | Sample Size | IMGT/HLA[1] Database | D |
|---|---|---|---|---|---|
| Papua New Guinea East New Britain Tolai | | 0.5650 | 48 | See | |
| Papua New Guinea Highland pop2 | | 0.3590 | 28 | See | |
| Papua New Guinea Highland | | 0.3590 | 94 | See | |
| Spain Andalusia Romani | | 0.3500 | 99 | See | |
| Sudan Central Region | 52.6 | 0.3100 | 97 | See | |
| Czech Republic Romani | 50.0 | 0.2928 | 34 | See | |
| China Shanghai pop 2 | | 0.2880 | 40 | See | |
| Sudan Mixed | 51.0 | 0.2880 | 200 | See | |
| Japan pop 2 | | 0.2820 | 916 | See | |
| South Korea pop 11 | | 0.2790 | 149 | See | |

**Figure S22.** Screenshot of DPB1*02:01 allele frequencies in modern human populations (high frequency top10 population data).

| Population | Phenotype Frequency (%) | Allele Frequency (in_decimals) | Sample Size | IMGT/ Data |
|---|---|---|---|---|
| Papua New Guinea Trobriand Island | | 0.9760 | 81 | Se |
| Taiwan Atayal pop 2 | | 0.8000 | 50 | Se |
| Taiwan Ami pop 2 | | 0.8000 | 50 | Se |
| Taiwan Rukai pop 2 | | 0.8000 | 50 | Se |
| Taiwan Bunun pop 2 | | 0.7800 | 50 | Se |
| Taiwan Saisiat pop 2 | | 0.7600 | 50 | Se |
| Taiwan Puyuma pop 2 | | 0.7600 | 50 | Se |
| Papua New Guinea Lowland Roro | | 0.7080 | 26 | Se |
| Samoa West | | 0.7040 | 22 | Se |
| Taiwan Aborigine pop 2 | | 0.6980 | 48 | Se |

**Figure S23.** Screenshot of DPB1*05:01 allele frequencies in modern human populations (high frequency top10 population data).

| Population | Phenotype Frequency (%) | Allele Frequency (in_decimals) | Sample Size |
|---|---|---|---|
| USA Arizona Pima pop2 | | 0.9410 | 18 |
| USA Arizona Gila River Amerindian | | 0.8960 | 492 |
| Colombia North Eastern Plains Sikuani | | 0.7780 | 27 |
| Colombia East Amazon Region Nukak | | 0.7250 | 20 |
| Russia Siberia Chukotka Peninsula Eskimo | | 0.6700 | 80 |
| Russia Siberia Polygus Evenk | | 0.6600 | 35 |
| Russia Siberia Eskimo | | 0.6440 | 70 |
| USA Alaska Yupik | | 0.6170 | 252 |
| Colombia West Waunana | | 0.6170 | 30 |
| Brazil Central Plateau Xavante | | 0.6150 | 74 |

**Figure S24.** Screenshot of DQB1*03:01 allele frequencies in modern human populations (high frequency top10 population data).

| Population | Phenotype Frequency (%) | Allele Frequency (in_decimals) | Sample Size |
|---|---|---|---|
| Colombia Sierra Nevada de Santa Marta Ijka pop 2 | | 0.6330 | 30 |
| Brazil Kaingang | | 0.5100 | 235 |
| Australia Kimberly Aborigine | | 0.4390 | 41 |
| USA New Mexico Canoncito Navajo | | 0.4250 | 42 |
| Colombia Sierra Nevada de Santa Marta Arhuaco | | 0.4150 | 107 |
| Mexico Chihuahua Tarahumara | | 0.3520 | 44 |
| Peru Titikaka Lake Uro | | 0.3340 | 105 |
| Peru Titikaka Lake Uros | | 0.3330 | 105 |
| Bolivia La Paz Aymaras | | 0.3100 | 87 |
| Bolivia Quechua | | 0.3043 | 69 |

**Figure S25.** Screenshot of DQB1*04:02 allele frequencies in modern human populations (high frequency top10 population data).

| Population | Phenotype Frequency (%) | Allele Frequency (in_decimals) | Sample Size |
| --- | --- | --- | --- |
| Colombia Sierra Nevada de Santa Marta Ijka pop 2 | | 0.6170 | 30 |
| Brazil Kaingang | | 0.4960 | 235 |
| USA New Mexico Canoncito Navajo | | 0.4380 | 42 |
| Colombia Sierra Nevada de Santa Marta Arhuaco | | 0.4150 | 107 |
| Peru Titikaka Lake Uros | | 0.3240 | 105 |
| Peru Titikaka Lake Uro | | 0.3190 | 105 |
| Mexico Mazahua | | 0.3100 | 65 |
| Bolivia La Paz Aymaras | | 0.3100 | 87 |
| Mexico Jaltepec Mazahuas | | 0.3100 | 50 |
| Mexico Sonora Seri | | 0.3030 | 34 |

**Figure S26.** Screenshot of DRB1*08:02 allele frequencies in modern human populations (high frequency top10 population data).

| Population | Phenotype Frequency (%) | Allele Frequency (in_decimals) | Sample Size |
|---|---|---|---|
| USA Arizona Pima pop2 | | 0.7940 | 18 |
| USA Arizona Gila River Amerindian | | 0.7460 | 492 |
| Colombia East Amazon Region Nukak | | 0.6500 | 20 |
| Colombia Waunana NA-DHS_20 (G) | 75.0 | 0.4750 | 20 |
| Russia Siberia NW of Sakhalin Island Nivkh | | 0.4200 | 32 |
| Colombia North Eastern Plains Sikuani | | 0.4070 | 27 |
| Colombia Sierra Nevada de Santa Marta Arsario | | 0.3500 | 18 |
| Russia Siberia Gvaysugi Udege | | 0.3500 | 25 |
| Canada British Columbia Athabaskan | | 0.3470 | 62 |
| Canada British Columbia Penutian | | 0.3460 | 26 |

**Figure S27.** Screenshot of DRB1*14:02 allele frequencies in modern human populations (high frequency top10 population data).

| Population | Phenotype Frequency (%) | Allele Frequency (in_decimals) | Sample Size |
|---|---|---|---|
| Russia Siberia Eskimo | | 0.3610 | 70 |
| Russia Siberia Chukotka Peninsula Chukchi | | 0.3000 | 59 |
| Russia Siberia Chukotka Peninsula Eskimo | | 0.2800 | 80 |
| Russia Siberia Chukchi | | 0.2540 | 71 |
| USA Alaska Yupik | | 0.2320 | 252 |
| Russia Siberia North East Kamchatka Koryak | | 0.2100 | 92 |
| Jordan Amman | | 0.1970 | 146 |
| Colombia Barranquilla | | 0.1860 | 188 |
| Denmark | | 0.1760 | 55 |
| Tunisia pop 2 | | 0.1700 | 111 |

**Figure S28.** Screenshot of DRB1*04:01 allele frequencies in modern human populations (high frequency top10 population data).

| Population | Phenotype Frequency (%) | Allele Frequency (in_decimals) | Sample Size |
|---|---|---|---|
| Papua New Guinea Kuru | | 0.3260 | 46 |
| Spain Pas Valley | | 0.3210 | 88 |
| Spain North Cabuerniga | | 0.3000 | 95 |
| Papua New Guinea Eastern Highlands Goroka Asaro | | 0.2980 | 57 |
| Papua New Guinea Ume | | 0.2890 | 90 |
| Papua New Guinea Woigi | | 0.2880 | 26 |
| Papua New Guinea Highland | | 0.2820 | 94 |
| Malaysia Kedah Baling Kensiu | | 0.2600 | 25 |
| Spain North Cantabria | | 0.2530 | 83 |
| Papua New Guinea Wonie | | 0.2160 | 51 |

**Figure S29.** Screenshot of DRB1*15:01 allele frequencies in modern human populations (high frequency top10 population data).

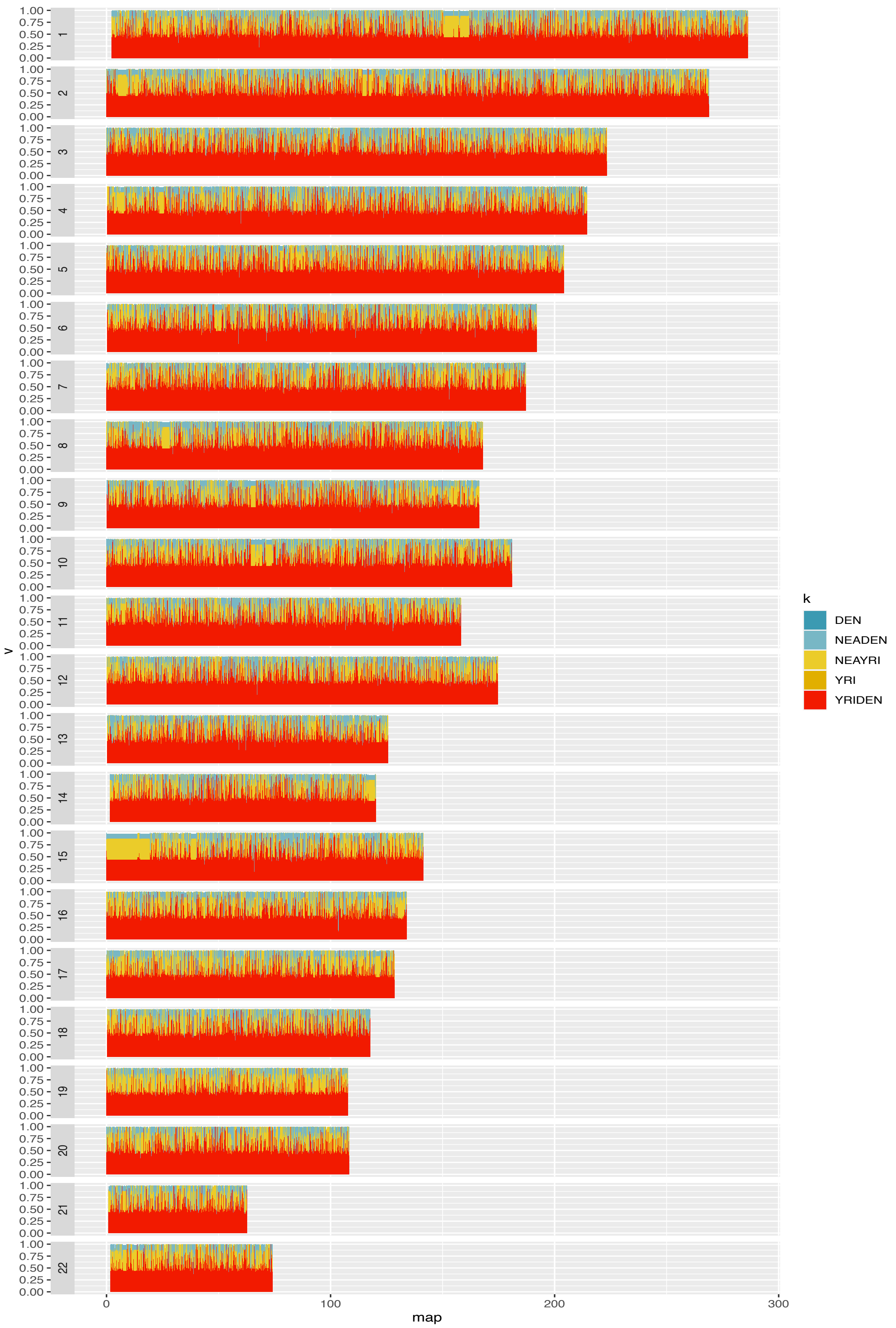


**Figure S30**. Inference of ancestry fragments in IY1 genome.
